# Gene networks with transcriptional bursting recapitulate rare transient coordinated expression states in cancer

**DOI:** 10.1101/704247

**Authors:** Lea Schuh, Michael Saint-Antoine, Eric Sanford, Benjamin L. Emert, Abhyudai Singh, Carsten Marr, Yogesh Goyal, Arjun Raj

**Affiliations:** Department of Bioengineering, University of Pennsylvania, Philadelphia, Pennsylvania 19104, USA; Institute of Computational Biology, Helmholtz Zentrum München, Neuherberg, 85764, Germany; Department of Mathematics, Technical University of Munich, Garching, 85748, Germany; Electrical and Computer Engineering, University of Delaware, Newark, Delaware 19716, USA

**Keywords:** stochasticity, network, gene expression, melanoma, drug resistance, non-genetic

## Abstract

**SUMMARY:** Non-genetic transcriptional variability at the single-cell level is a potential mechanism for therapy resistance in melanoma. Specifically, rare subpopulations of melanoma cells occupy a transient pre-resistant state characterized by coordinated high expression of several genes. Importantly, these rare cells are able to survive drug treatment and develop resistance. How might these extremely rare states arise and disappear within the population? It is unclear whether the canonical stochastic models of probabilistic transcriptional pulsing can explain this behavior, or if it requires special, hitherto unidentified molecular mechanisms. Here we use mathematical modeling to show that a minimal network comprising of transcriptional bursting and interactions between genes can give rise to rare coordinated high states. We next show that although these states occur across networks of different sizes, they depend strongly on three (out of seven) model parameters and require network connectivity to be ≤ 6. Interestingly, we find that while entry into the rare coordinated high state is initiated by a long transcriptional burst that also triggers entry of other genes, the exit from it occurs through the independent inactivation of individual genes. Finally, our model predicts that increased network connectivity can lead to transcriptionally stable states, which we verify using network inference analysis of experimental data. In sum, we demonstrate that established principles of gene regulation are sufficient to describe this new class of rare cell variability and argue for its general existence in other biological contexts.

## INTRODUCTION

Both natural and synthetic systems exhibit rare and large deviations from their typical behavior, ranging from the occurrence of hurricanes to dramatic drops in the stock markets (Taleb, 2007). Biology is also replete with examples of rare deviations in organismal or cellular behavior. The most prototypical example, pertaining to individuals within a population, is evolution, where rarely occurring mutations can lead to organisms with different traits. Cancer is another such example (at the level of cells within a population), in which rare cells acquire mutations that drive uncontrolled cellular proliferation. In the vast majority of examples of rare biological behavior examined to date, the driver of the rare behavior is thought to be genetic, i.e. a result of mutation(s) in the DNA. However, recent studies have suggested that rare and large deviations in single cells, such as in cancer, can also be driven by non-genetic sources, even in clonal, genetically homogeneous cells grown in identical conditions (Fallahi-Sichani et al., 2017; Gupta et al., 2011; Pisco and Huang, 2015; Shaffer et al., 2017; Sharma et al., 2018, 2010; Spencer et al., 2009; Su et al., 2017). Importantly, cells exhibiting these non-genetic deviations are resistant to anti-cancer drugs (e.g., Ras pathway inhibitors) and may lead to relapse in patients.

In the case of melanoma, a small fraction (∼1 in 3000) of cells are pre-resistant, meaning they are able to survive targeted drug therapy, resulting in their uncontrolled cellular proliferation. These rare pre-resistant cells are marked by transient and coordinated high expression of several marker genes. Notably, the expression levels of marker genes in individual cells are not normally distributed, instead showing a heavy-tailed, subexponential distribution. These rare cells in the tails, which transiently arise and disappear in the population by switching their gene expression state (**Figure 1A**), are much more likely to develop resistance to targeted therapies. Thus, a larger fraction of cells express very high levels of multiple marker genes compared to the bulk population than one would expect for a normal distribution (**Figure S1A**). Importantly, this type of single cell variability, which is characterized by rare and coordinated large deviations in the expression of multiple genes, is conceptually distinct from the classical “noise” models of non-genetic single cell variability using probabilistic models of gene regulation (Antolovic et al., 2017; Chen and Larson, 2016; Corrigan et al., 2016; Golding et al., 2005; Raj and van Oudenaarden, 2008; So et al., 2011; Symmons and Raj, 2016; Thattai and van Oudenaarden, 2001). Specifically, the classical models have largely described the variability that results in relatively normally distributed counts of mRNAs of a given gene per cell. However, in this classical context, most of the cells in the tail of this distribution are not very different from the bulk of the population, a scenario distinct from that described above for melanoma cells (**Figure S1A**). This presents an opportunity to investigate the origins of these rare, transient, and coordinated high gene expression states (from now on referred to as “rare coordinated high states”).

**Figure 1.**
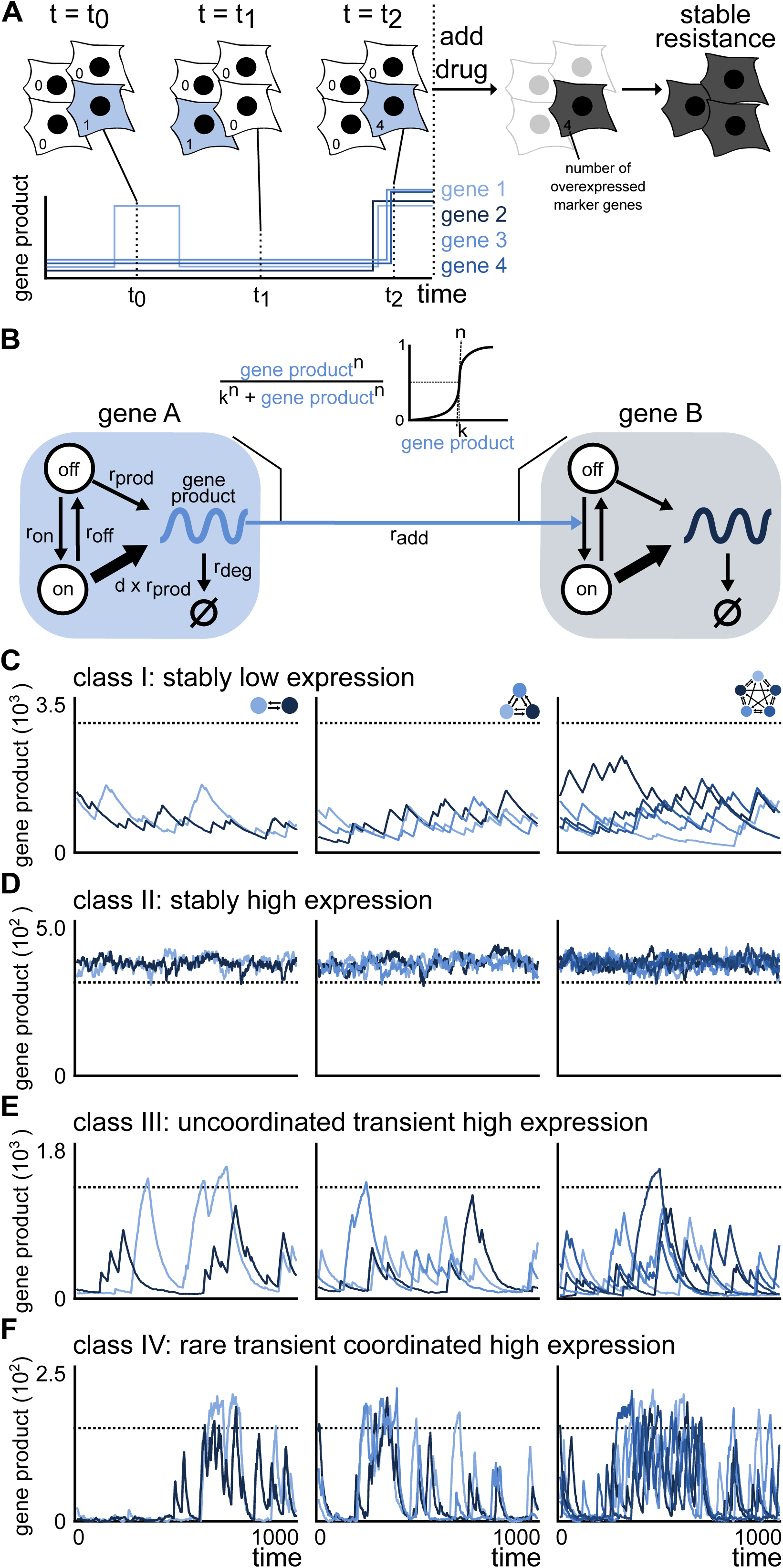
A transcriptional bursting-based stochastic network model is able to mimic the rare coordinated high states observed in melanoma. (A) Drug naive melanoma cells exist in low (white cells) as well as rare, transient, and coordinated high (blue cells) expression state. Cells in the rare transient coordinated high expression state characterize the pre-resistant state observed in melanoma. A schematic of the corresponding expression pattern is shown in the panel below. The cells in a high expression state are more likely to survive and acquire resistance upon drug administration. (B) Schematic of the transcriptional bursting-based stochastic network model for two nodes. DNA is either in an inactive (off) or active (on) state. Transitions take place with rates r_on_ and r_off_, where mRNA is synthesized with rates r_pod_ and d*r_prod_, respectively. mRNA degrades with rate r_deg_. Gene regulation is modeled by a Hill function, where the gene expression of the regulating gene A increases the activation of the regulated gene B. (C-F) Depending on the network architecture and the parameters of the gene expression model, we observe stably low expression (C), stably high expression (D), uncoordinated transient high expression (E) and rare transient coordinated high expression (F). See also Figure S1.

Might a stochastic system of interacting genes inside the cell facilitate transition in and out of the rare high expression state? Previous efforts using the canonical stochastic models of gene regulation to study large deviations have been limited to characterizing the ensuing gene expression distributions in isolated, single gene reaction systems (Ham et al., 2019; Horowitz and Kulkarni, 2017; Iyer-Biswas et al., 2009; Stinchcombe et al., 2012). It is not clear if such models can explain the rare, transient, and coordinated large deviations observed in gene expression profiles of multiple genes. One hypothesis is that within the canonical modeling framework, only a rare set of unique (and perhaps complex) network architectures can facilitate reversible transitions into the rare coordinated high states. Alternatively, relatively generic gene regulatory networks may be capable of producing such behaviors, suggesting that a large ensemble of such networks may admit rare-cell formation. Both of these scenarios have different implications—for instance, the latter hypothesis suggests that this behavior could be more common in biological systems than hitherto appreciated. The alternatives described above can also be posed in terms of the nature of model parameters—whether the set of values that give rise to rare coordinated high states are constrained to lie within a narrow window of parameter space or whether such behavior may occur across broad swaths of parameter space. Yet another possibility is that standard stochastic gene expression models fail to produce rare coordinated high states entirely, no matter what combinations of network architectures and parameters are used. In that case, one may argue that this reversible transition into the rare coordinated high state is driven by highly specialized processes (e.g. initiated by a master regulator) or other unknown mechanisms.

Here we describe a mathematical framework to test the hypotheses proposed above for the appearance and disappearance of rare and coordinated high expression cellular states. Recent studies from our lab suggest that no particular molecular pathway is solely responsible for the formation of these rare cells (Shaffer et al., 2018; Torre et al., 2019). Specifically, in these rare cells, a sequencing and imaging based scheme identified a collection of marker genes, which are targets of multiple signaling pathways. This implies that instead of a single signaling pathway leading to the observed behavior, a network of interacting genes appears to be responsible. Accordingly, our approach used network modeling to see whether genes interacting within this framework were capable of producing transitions to high expression states. We systematically formulated and simulated networks of increasing size and complexity defined by a broad range for all free parameters (**Methods**, section Network architectures & section Parameters).

Our computational screens on more than 96 million simulated cells reveal that many networks with stochastic interactions between genes are capable of producing rare coordinated high states. Critically, transcriptional bursting, a ubiquitous phenomenon in which genes flip between transcriptionally active and inactive state, is necessary for our model to produce these rare coordinated high states. Subsequent quantitative analysis shows that rare coordinated high states occur across networks of all sizes, but that they (i) depend on three (out of seven) model parameters and (ii) require network connectivity to be ≤ 6. The transition into the rare coordinated high state is initiated by a long transcriptional burst, which, in turn, triggers the entry of subsequent genes into the rare coordinated high state. In contrast, the transition out of rare coordinated high state is independent of the duration of transcriptional burst, rather it happens through the independent inactivation of individual genes. We also confirm some model predictions using experimental gene expression data taken from melanoma cell lines. Together, this demonstrates that the standard model of stochastic gene regulation with transcriptional bursting is capable of producing rare, transient, and coordinated high expression states.

## RESULTS

### Framework selection

#### Identifying the minimal network model generating rare coordinated high states

We focused on a network-based mathematical framework that models cell-intrinsic biochemical interactions and wondered what would be the minimal set of biochemical reactions that constitutes it. Since network models without gene activation (i.e. constitutive mode of gene expression) were not able to produce rare, transient, and coordinated high expression states (see **Supplementary information**; **Figure S1B-D**; **Methods**, section Model 1), we use a telegraph model as the building block of our framework. In this model, we incorporate transcriptional bursting at each node, a phenomenon in which genes flip reversibly between transcriptionally active and inactive state regulated by the binding of a transcription factor(s) (**Figure 1B**; **Methods**, section Model 2).

In terms of chemical reactions, a gene can reversibly switch between an active (r_on_) and inactive state (r_off_), where binding of the transcription factor at a gene locus controls the effective rate of gene production (**Figure 1B**). Specifically, when inactive (or unbound), the gene is transcribed as a Poisson process at a low basal rate (r_prod_); when active, this rate becomes higher (*d x r*_*prod*_, where *d* > *1*). We modeled degradation of the gene product as a Poisson process with degradation rate r_deg_. The inter-node interaction parameter, r_add_, has a Hill-function-based dependency on the gene product amount (Hill coefficient n) of the respective regulating node to account for the multistep nature of the interaction (**Figure 1B**). All chemical reactions, propensities, and model parameters are presented in **Methods**. We used Gillespie’s Stochastic Simulation Algorithm (Gillespie, 1977) to systematically simulate networks of various sizes and architectures across a broad range of parameters (**Methods**, section Network architectures & section Parameters).

We started with the simplest subset of the millions of possible networks (**Figure S1E**) to see if we could find the rare transient and coordinated high gene expression states. We limited our study to networks that are symmetric, i.e., networks without a hierarchical structure (**Methods**, section Network Architectures). This simplification is partially supported by the experimental observation that there doesn’t seem to be a clear correlation between the genes that are highly expressed in the transient rare state in melanoma (**Figure S1F**) (Shaffer et al., 2017, 2018). Symmetric models also allow for comparisons of parameters between networks of different sizes. Additionally, we excluded networks that are compositions of independent subnetworks (non weakly-connected networks) and networks that can be formed by renaming other networks (isomorphic networks) (**Methods**, section Network Architectures). With these restrictions, we avoid analyzing same network architectures several times -- either as independent subnetworks in larger architectures -- and/or as reordered forms of other architectures. With these operations, we also reduce the testable space of unique network architectures by several orders of magnitude (**Figure S1E**).

#### Characterization of the transcriptional bursting network model

When the genes are organized in the system described above and simulated over long intervals, our stochastic model produced a range of temporal profiles for gene products (**Figure 1C-F** and **Figure S2A**). Importantly, this model was able to faithfully capture the qualitative features of the experimental data, i.e., rare, transient, and coordinated high expression states (**Figure 1F**). We defined a set of rules to screen for the occurrence of different classes of states (**Figure 1C-F** and **Figure S2A**); these include stably low expression (class I), stably high expression (class II), uncorrelated transient high expression (class III), and rare transient coordinated high expression (class IV) (see **Methods**, section Simulation classes), and used a heuristic approach to distinguish between these different classes. In particular, we defined the rare, transient, and simultaneous production of multiple gene products as equivalent to the experimentally observed rare coordinated high state when half or more of the genes in a given network are transiently in the high expression state (blue cell in **Figure 1A**) at the same time. For a detailed description of the rules and quantitative metrics used to define class IV, see **Figure S3** and **Methods**, section Simulation classes.

To better compare our computational results with the experimental data from static RNA FISH images, we took snapshots of gene products at randomly selected time points in our simulations and noted the number of co-occurring gene products as well as their counts for each time point (**Figure 2A**). This exercise, justified by ergodic theory, allows us to represent the static states of an ensemble of (computational) cells. For example, in a particular 8-node network, we found that the distribution qualitatively captures the experimental observations where most cells do not exhibit high expression states, while some cells are in a high-state for one or more genes **(Figure 2B)**. Similarly, when we selected a gene and plotted its product count for randomly chosen time points, we observe a heavy-tailed distribution (**Figure 2C**), similar to the experimental observations **(Figure 2C)**. Furthermore, these observations, while shown for a particular 8-node network, hold true for simulations of other 8-node network architectures as well as the networks of other sizes (**Figure S2B**). Note that the distributions of gene product counts for each gene are qualitatively similar because of the symmetric nature of the networks (**Figure S2C**). The experimental data in melanoma cells for gene expression counts display different degrees of skewness of the distribution for different genes. This can likely be recapitulated by introducing asymmetries in the networks. To test this, we randomly introduced asymmetry either in the network architecture or the model parameters. The two asymmetric networks we tested were both able to produce rare, transient and coordinated high expression states (**Figure S4A-S4F**). Importantly, the distributions of gene expression counts for various genes displayed different levels of heavy-tails, as also observed in the experimental data (**Figure S4G**). Together, the transcriptional bursting-based network model is able to produce states which recapitulate key aspects of rare coordinated high states observed in melanoma.

**Figure 2.**
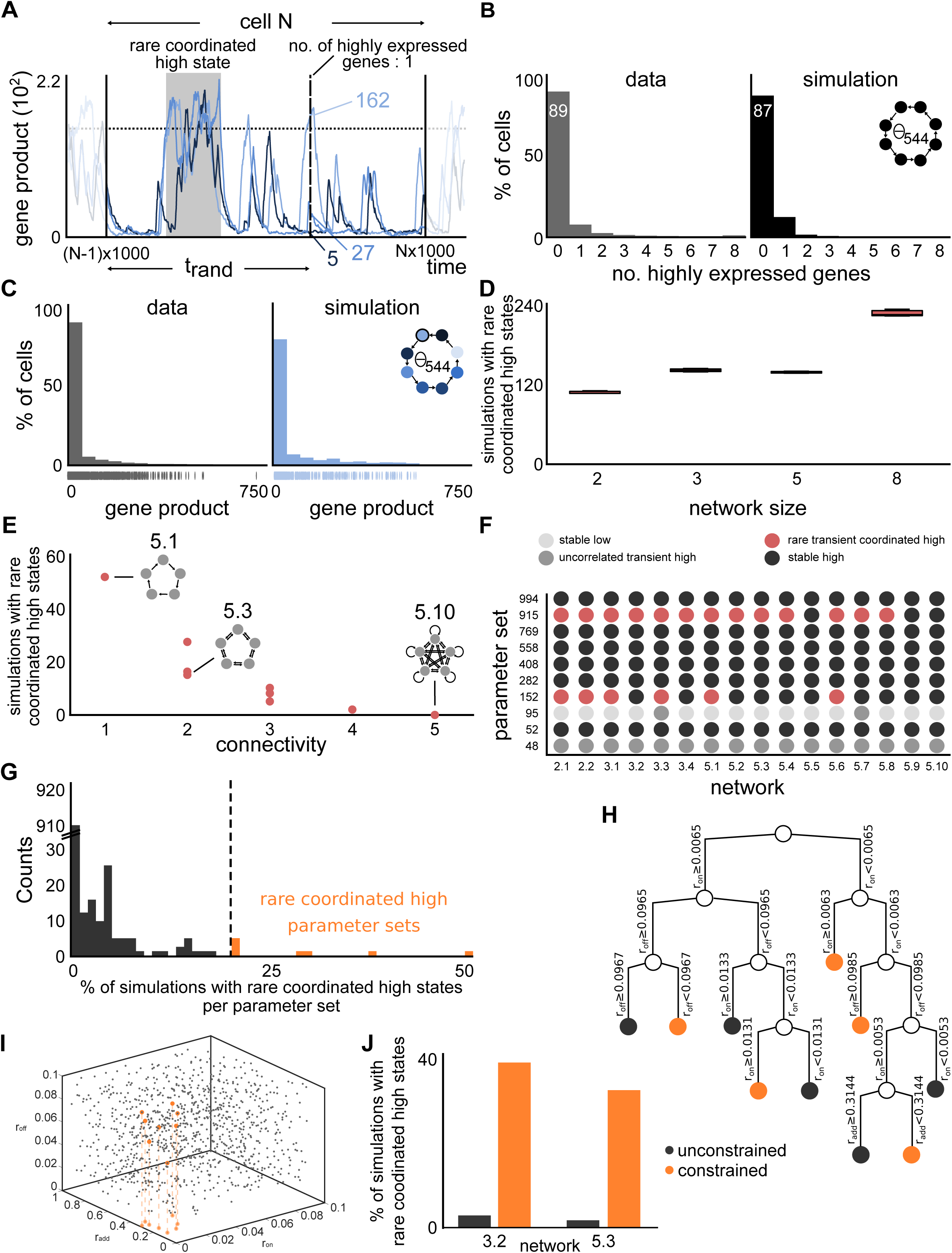
Transcriptional bursting-based stochastic network model simulations show similar behavior at the population level as the melanoma cells. (A) Frame of simulation showing rare coordinated high state (shaded area). The 1000000 time unit simulation is split into frames of 1000 time units to create a simulated cell population (shown for cell N). For a randomly determined time-point t_rand_, the number of highly expressed genes and the gene count per gene per cell are evaluated. (B,C) The simulated number of simultaneously highly expressed genes and expression distribution are qualitatively similar to experimental data from a drug naive melanoma population. (data from (Shaffer et al., 2017)) (D) Rare coordinated high states occur ubiquitously across networks of all analyzed network sizes. (E) Increasing connectivity within all networks of size 5 leads to a decrease in the number of simulations with rare coordinated high states. (F) Simulations of a particular parameter set across different network architectures and sizes show largely the same class of gene expression profiles. (G) The eight rare coordinated high parameter sets give rise to rare coordinated high states more frequently than others for all 96 networks and cluster at the tail of the histogram. (H) Decision tree optimization of resulting parameters reveal that only three out of seven parameters, r_on_, r_off_, and r_add_, show a strong correlation with the rare coordinated high state producing parameter sets. (I) Three dimensional representation of all tested 1000 parameter sets for r_on_, r_off_, and r_add_ show that the rare coordinated high parameters are narrowly constrained in the 3D space (orange). (J) The constrained subregion defined by the rare coordinated high parameter sets heavily favors the formation of rare coordinated high states. See also Figure S2, Figure S3, Figure S4, and Figure S5.

### Rare coordinated high states depend on network connectivity and model parameters

What kind of network architectures and model parameters might facilitate the occurrence of rare coordinated high states? For the simulations that produced rare coordinated high states, we extracted and quantitatively analyzed the corresponding network architectures and parameter values. We found that the rare coordinated high states occurred ubiquitously in networks with different numbers of nodes (analysed for up to 8 nodes) (**Figure 2D**). This suggests that even larger networks (≥ 8 nodes) will also display rare coordinated high states. Next, we wondered if the occurrence of rare coordinated high states depend on the network connectivity, defined as the number of ingoing edges for any node in the network. Indeed, we found that, within a particular network size, the ability to produce rare coordinated high states decreases dramatically (and monotonically) with increasing network connectivity (**Figure 2E and Figure S2F**), with a peak frequency for a connectivity of one. The rare coordinated high states cease to exist for a network connectivity > 6 (**Figure S2G**).

Our model has seven free parameters and we asked whether the occurrence of the rare coordinated high states is also controlled by parameter combinations. To test this, we ran the simulation for 1000 parameter combinations sampled from a broad range for each parameter using a Latin Hypercube Sampling algorithm across networks of different sizes and architectures (**Supplementary Information** ParSetsAnalysis.xlsx**; Methods**, section Parameters). Occurence of different classes of temporal gene product profiles (**Figure 1C-F**) across different network sizes and connectivities depend on the parameter sets (**Figure 2F**). Specifically, if a parameter set gave a specific expression profile (e.g. rare coordinated high or stably high) for one network, it displayed a higher propensity to display the same profile for other networks as well (**Figure 2F** and **Figure S2H**). This implies that parameters indeed play a major role in the occurrence of rare coordinated high states. We therefore measured the percentage of simulations per parameter set that gave rise to the rare coordinated high states (**Figure 2G**). Out of the 1000 parameter sets, eight parameter sets clustered together at the tail-end of the distribution (orange, **Figure 2G**), meaning they generated simulations with frequent occurrence of rare coordinated high states, at least 20% of all networks tested (**Figure 2G**). Furthermore, these 8 parameter sets robustly generated rare behaviors across all network sizes and architectures even when we subsampled networks of specific properties, e.g. fixed size or connectivity (**Figure S5A** and **Figure S5B**). Intrigued by this observation, we wondered if these eight parameter sets have any special or distinguishing features compared to the remaining 992 parameter sets.

We used a decision tree algorithm-based (Breiman et al., 1984) approach (see **Methods**, section Decision tree optimization and generalized linear models) to identify the differentiating features of these parameter sets from the rest, and confirmed our results with additional simulations. Our analysis revealed that only three (r_on_, r_off_, and r_add_) of the seven free parameters showed a strong correlation with the rare coordinated high state producing parameter sets (**Figure 2H**). We validated these findings with complementary analysis using generalized linear models (**Methods**, section Decision tree optimization and generalized linear models) where we found precisely these three specific parameters to be critical to produce the rare coordinated high states with high statistical significance (p values: r_on_ = 0.003; r_off_ = 0.005; r_add_ = 0.014) (**Figure S5C**). These observations became readily evident when we plotted all the 1000 simulated parameters for r_on_, r_off_, and r_add_ together and found the rare coordinated high state producing parameters to occupy a narrow region of the parameter space (**Figure 2I** and **Figure S5D**). All of these parameters are related to transcriptional bursting and inter-gene(node) regulation. Two of these parameters, r_on_ and r_off_, define the active and inactive state of the gene respectively.

The third parameter is the gene activation rate, r_add_, which corresponds to the positive regulation of transcriptional bursting rate of a gene by the gene product of another interacting gene. Interestingly, too high value (> 0.31) of r_add_ results in the disappearance of rare coordinated high states, as does a complete absence (r_add_ = 0) of this term (**Figure S5E-S5G**). To confirm that these three parameters (r_on_, r_off_, and r_add_) and their corresponding range of values are indeed critical to producing rare behaviors, we sampled new 1,000 parameter sets from the region constrained for these three parameters (**Methods**) and ran simulations for two test networks, a 3-node and a 5-node network. We found that the frequency of pre-resistant states for the constrained region is ∼14-fold and ∼21-fold higher than that for the original parameter set for the 3-node and the 5-node network, respectively (**Figure 2J**). Together, network connectivity and three model parameters are primarily regulators of the propensity to display rare, transient, and coordinated high expression states.

### Distinct mechanisms regulate the transition into and out of rare coordinated high states

We have identified the network architectures and parameter sets for which our model exhibits rare coordinated high states. Next, we wondered if we could dissect the features of the model that facilitate this behavior. Specifically, we wanted to know what factors trigger the entry into the rare coordinated high states and what factors trigger the escape from it. We began by analyzing the length of transcriptional bursts (burst duration), since including transcriptional bursting parameters was critical for the model to display the rare coordinated high state. Accordingly, we measured transcriptional burst durations in four regions of each simulation: low expression state (baseline time-region), entry into the high expression state (entry time-point), the high expression state (high time-region), and exit from the high expression state (exit time-region) (**Figure 3A, Methods**, section Entry and Exit mechanisms).

**Figure 3.**
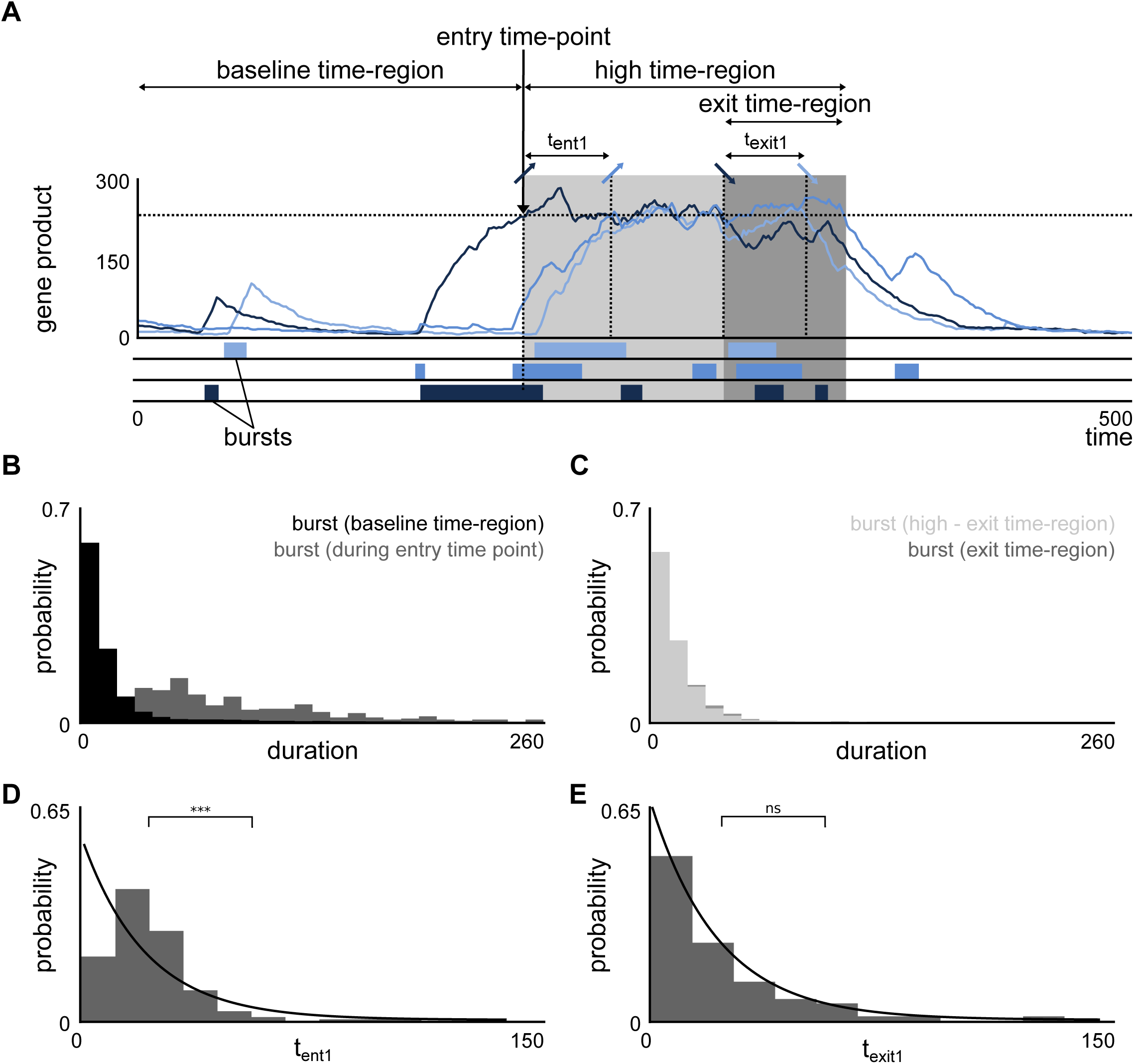
Rare coordinated high state is initiated by a long transcriptional burst, while its termination is a random process. (A) An exemplary high region, with a baseline time-region, entry time-point, high time-region and an exit time-region. The time intervals for an additional gene to enter and exit the high region are marked by t_ent_ and t_exit_, respectively. (B) The bursts during entry time-points are significantly longer than bursts not in a high time-region. (C) There is no statistical significant difference between the distributions underlying the duration of bursts in the high time-region and the exit time-region. (D,E) The time intervals between genes entering and exiting the high time-region are distributed differently, as demonstrated by two representative simulations. While the time intervals for entering the high time-region are not exponentially distributed (E) (and hence not random), the time intervals for exiting the high time-region are exponentially distributed (F). See also Figure S6.

We found that for any gene in the network, the transcriptional burst duration right before and during the entry into a rare coordinated high state was significantly higher than that in the baseline time-region (i.e., regular bursting kinetics) (**Figure 3B** and **Figure S6A**). In the example shown in **Figure 3B**, the average time of transcriptional burst at the entry time-point is 84.82 (time units) as compared to only 15.08 (time units) in the baseline time-region. This implies that prolonged transcriptional bursts play a role in driving the cell to a coordinated high expression state. Conversely, we asked if the opposite is true at the exit time-region, such that transcriptional bursts for the exit time-region are shorter than for the high time-region. We found no difference in the distributions of burst durations between the high and the exit time-regions (**Figure 3C** and **Figure S6A**). This suggests that the exit from high expression state occurs independently of the burst durations. Therefore, unlike the entry into the high time-region, the exit from it is not dependent on the transcriptional burst duration.

We also wondered if the entry into the high expression state of one gene influences the entry of other genes, or that the genes enter the high expression state independent of each other. We reasoned that if the time duration between two successive genes (t_ent_, **Figure 3A**) entering the high expression state is exponentially distributed, it would imply that the genes enter the high expression state independent of each other. Instead, we found that the distributions of entry time intervals rejected the null-hypothesis of the Lilliefors’ test for most of the simulations (84%), meaning they are not exponentially distributed (**Figure 3D**). The remaining 16% of cases were found to be largely falsely identified as exponentially distributed due to limited data (see a representative example in **Figure S6B)**. This suggests that long transcriptional bursts at genes seem to reinforce one another through the interactions between network nodes (as defined by parameter r_add_, **Figure 1B**) and organize into a “super-burst”, resulting in the high expression of multiple genes for a sustained period. Similarly, we tested if the exit for successive genes from the high expression state occurs independent of each other. Interestingly, contrary to the situation during the entry into the high expression state, many distributions of exit time intervals satisfied the null-hypothesis of the Lilliefors’ test, implying they are indistinguishable from exponential distributions (**Figure 3E**). The simulations that did not satisfy the stringent Lilliefors’ test mainly appear to be exponentially distributed nevertheless; a representative example is shown in **Figure S6C**. Together, the entry into and exit from the rare coordinated high state occur through fundamentally different mechanisms — the entry of one gene into the high expression state affects the entry of next gene, while they exit from it largely independently of each other.

### Increasing network connectivity leads to transcriptionally stable states

So far, we have used our model to understand the potential origins of rare pre-resistant states in melanoma cells that are not exposed to drug. Upon treatment with anti-cancer drugs, the transient pre-resistant drug naive cells reprogram and acquire resistance resulting in their uncontrolled proliferation. The resistant cells are characterized by the stabilization of the high expression of the marker genes which were transiently high in the drug naive pre-resistant cells (**Figure 4A**) (Shaffer et al., 2017). Studies using network inference of gene expression data have suggested that the genetic network architectures undergo significant rearrangements upon cellular transitions or reprogramming (Moignard et al., 2015; Schlauch et al., 2017). We wondered if our network model can explain how might the transient high expression in drug naive cells become permanent upon treatment with anti-cancer drugs. Our modeling framework produces a range of gene expression profiles, depending on the network properties and model parameters (**Figure 1C-F**). Increasing the network connectivity (for fixed parameter values) is one way to shift from a transient coordinated high expression state to stably high expression state (**Figure 4B-E**). In particular, the model predicts that in networks with high connectivity, cells can transition into the high expression state but lose the ability to come out of it (**Figure S7A** and **S7B**). As an example, for a fixed network size (five) and associated parameters, increasing the network connectivity from one to five (amounting to 3.6 fold increase in total connections) resulted in a shift from transiently to stably high gene expression states (**Figure 4D** and **Figure 4E**). This is also reflected by the bimodal distribution of the abundances of genes product counts for randomly chosen timepoints in the highly connected network (**Figure 4F** and **Figure 4G**), where genes stay permanently in the high state once they leave the low expression state. These results mimic the experimentally measured gene expression states of the drug-induced reprogrammed melanoma cells.

**Figure 4.**
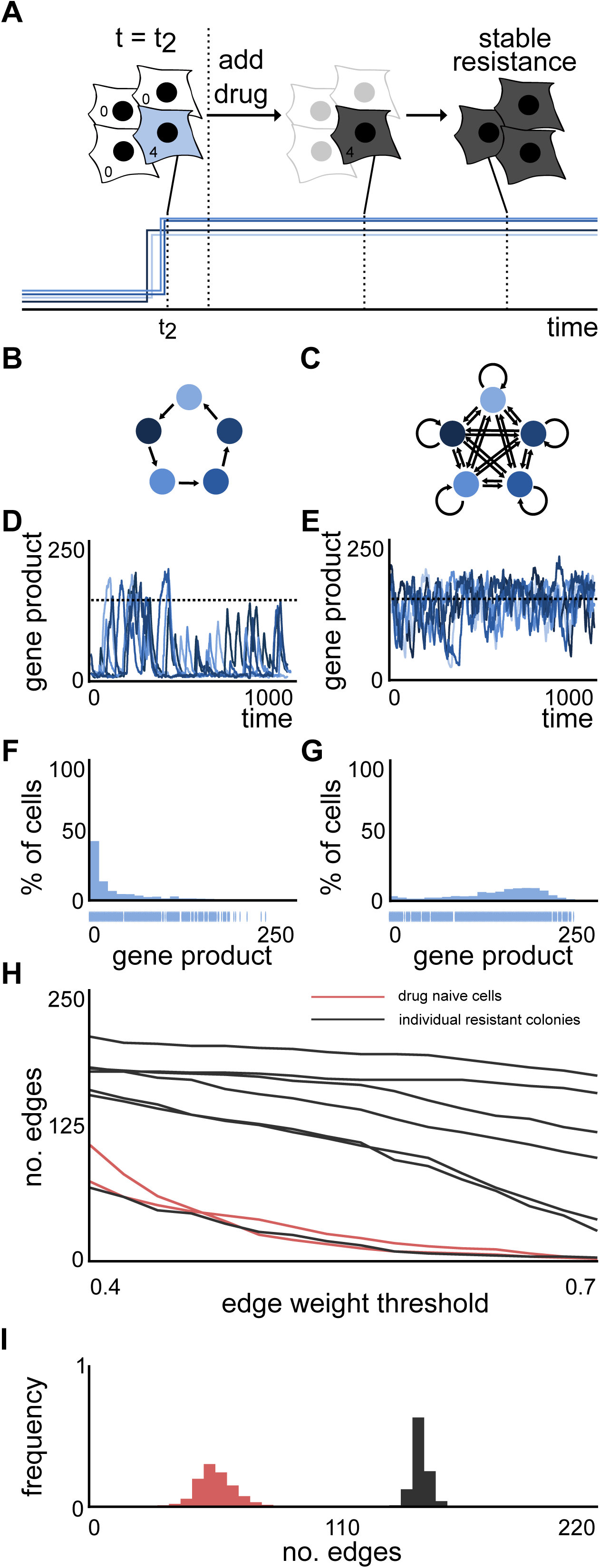
Increased connectivity of a network leads to stable high expression which is also observed in emerging resistant colonies post-drug treatment. (A) Upon drug treatment, the surviving cells acquire stable resistance. A schematic gene expression pattern is shown below. (B,C) Network architectures of size 5 with low (B) (1) and high (C) (5) connectivity and corresponding (D,E) simulations. (F,G) The gene expression distribution underlying the simulation of the highly connected network (G) does not exhibit heavy-tails while the simulation of the lowly connected network (F) exhibits heavy-tails. (H) Network inference analysis shows that 6 out of 7 resistant colonies have higher connectivity in comparison to two biological replicates of drug naive cells for many edge weight thresholds. (I) Distribution of number of edges for the drug naive cells (red) is lower than an exemplary resistant colony (black) when the network inference analysis is run 1000 times on bootstrapped data. See also Figure S7 and Figure S8.

To test if this computational prediction holds true in melanoma, we performed network inference using Φ-mixing coefficient-based (Ibragimov, 1962) Phixer algorithm (Singh et al., 2012) on the experimental data (**Methods**, section Comparative Network Inference). Specifically, we used the Phixer algorithm on the mRNA counts obtained from fluorescent in situ hybridization (FISH) imaging data of marker genes in drug naive cells and the resistant colonies that emerge post-drug treatment to infer the underlying network architecture. Consistent with our model prediction, we found that the number of edge connections (for a range of edge weight thresholds) between marker genes increased more than two-fold for 6/7 resistant colonies compared to the drug-naive cells (**Figure 4H**). To control for biases from subsampling of the experimental data and the nature of phixer algorithm itself (see **Methods**, section Comparative Network Inference), we ran the entire network inference analysis 1000 times. Again, in all 1000 runs, we saw a higher number of total edges for 6/7 resistant colonies compared to the drug-naive cells (**Figure 4I, Figure S7C** and **Figure S8**).

Besides the dependence on network architures, our framework predicts that for a given network architecture, stronger interactions between nodes (defined by the interaction parameter r_add_) can also result in stable gene expression profiles (**Figure S5E-S5G**). While measuring the strength of interactions between molecules is experimentally challenging, it is likely that reprogramming results from a combination of increased edge connectivity as well as the enhanced interactions (given by parameter r_add_) between existing edges. Biologically, this translates into stronger and increased number of interactions between genes and associated transcription factors during reprogramming. Together, network inference of our experimental data is consistent with model findings about the cellular progression from a transient high expression state to a stable state.

## DISCUSSION

We developed a computational framework to model rare cell behaviors in the context of melanoma where a rare subpopulation of cells displays transient and coordinated high gene expression states. We found that a relatively parsimonious model consisting of transcriptional bursting and stochastic interactions between genes in a network is capable of producing rare behaviors that fully mimic the experimental observations. To systematically investigate their origins, we screened networks of increasing sizes and connectivities for a broad range of parameter values. Our study revealed that their occurrence is dependent on 3/7 model parameters and requires the network connectivity to be ≤ 6. Furthermore, we showed that the mechanisms that lead to the transition into- and out of- the rare coordinated high expression state are fundamentally different from each other. Collectively, our framework provides an excellent basis for further mechanistic and quantitative studies of the origins of rare, transient, and coordinated high expression states.

Given the relative generality of the scenarios that produce rare behaviors (Shaffer et al., 2018), our model predicts that every cell type is capable of entering the transient and coordinated high gene expression state. In the case of melanoma cells, this transient state is characterized by an increased ability to survive drug therapy leading to uncontrolled proliferation of the resulting resistant cells. It is possible that these rare transient behaviors exist across many cell types and have a variety of phenotypic consequences. Our model also makes two key predictions regarding transitions into and out of rare coordinated high states. The first is that prolonged transcriptional bursts drive entry into the high expression state while exit from it is independent of the burst duration. The second is that genes entering the high expression state promote the entry of subsequent genes, whereas genes exiting the high expression state do so independently of each other. Both these predictions can be readily tested experimentally by simultaneous visualization of transcriptional bursting and mRNA counts using live cell (e.g. by using RNA-binding fluorescent proteins) or fixed cell (intron and exon FISH) imaging approaches (Bartman et al., 2016; Rodriguez et al., 2019).

Additionally, we showed that increasing the network connectivity is one way to reach a drug-induced reprogrammed state, a prediction we verified by performing network inference analysis on the experimental data. Our model proposes that there are many plausible ways to transition from networks that produce transient high expression states to stable high expression states. For example, this transition could be facilitated by different amounts of increases in connectivities between nodes (genes) and/or changes in parameter values of the gene expression model. Furthermore, it is possible that these changes may take place only for a subset of nodes and edges within the network. These computational scenarios suggest that there could be significant heterogeneity in the stable expression levels of network genes in the resistant colonies emerging even from clonal population of drug naive cells. This possibility can be tested experimentally by isolating individual colonies and profiling them for molecular markers to identify the paths. Identification of dominant paths has relevance for rational targeted drug therapy design. Therefore, in addition to modeling rare-behaviors, our framework can be adapted for investigating the plasticity and reprogramming paradigm in cancer.

One limitation of our model is that we have performed quantitative analysis only on symmetric networks with positive interactions between nodes. It is likely that our findings hold more generally for asymmetric networks, as partially demonstrated for two cases of randomly selected asymmetric networks (**Figure S4A-S4G**). Inhibitory interactions between nodes is a separate and perhaps more interesting point. In principle, the model can be adapted to include inhibitory interactions. These inhibitory interactions may lead to non-monotonic effects of network connectivity on the occurrence of rare states, as positive and negative interactions can compete in non-linear ways. Inclusion of these interactions might also make the exit of genes from the high expression state dependent on one another, which occurs independently in our current model.

While we have focused on rare, transient, and coordinated high expression states in melanoma, our study provides conceptual insights into other biological contexts such as stem cell reprogramming. Particularly, there is increasing evidence to suggest that stem cell reprogramming to desired cellular states proceeds *via* non-genetic mechanisms in a very rare subset of cells (Hanna et al., 2009; Pour et al., 2015; Takahashi and Yamanaka, 2016). Our model may explain the origins and transient nature of this type of rare cell variability. In sum, we have established the plausibility that a relatively parsimonious model comprising of transcriptional bursting and stochastic interactions of genes organized within a network can give rise to a new class of biological heterogeneities. In light of this, we believe that established principles of transcription and gene expression dynamics may be sufficient to explain the extreme heterogeneities that are being reported increasingly in a variety of biological contexts.

## Supporting information

ParSetsAnalysis.xlsx

PhixerData.xlsx

## SUPPLEMENTAL INFORMATION

Supplemental Information includes 9 figures and 2 tables.

## ACKNOWLEDGEMENTS

We thank the Raj lab members, especially Ian Mellis and Amy Azaria, for scientific discussions and comments on the manuscript. We also thank Ravi Radhakrishnan and Alok Ghosh for helpful discussion during the initial stages of this project. We thank Cesar A Vargas-Garcia for his help during the initial discussions on network inference. A.R. acknowledges NIH/NCI PSOC award number U54 CA193417, NSF CAREER 1350601, P30 CA016520, SPORE P50 CA174523, NIH U01 CA227550, NIH 4DN U01 HL129998, NIH Center for Photogenomics (RM1 HG007743), and the Tara Miller Foundation. C.M. acknowledges support from the Deutsche Forschungsgemeinschaft DFG through the SFB 1243. A.S. acknowledges support from the NIH grant 5R01GM124446-02. L.S. would like to acknowledge the support of the PROMOS fellowship of the DAAD, Germany. Y.G. would like to acknowledge the Schmidt Science Fellows in partnership with the Rhodes Trust. Y.G. is a fellow of The Jane Coffin Childs Memorial Fund for Medical Research and this investigation has been aided by a grant from The Jane Coffin Childs Memorial Fund for Medical Research.

## AUTHOR CONTRIBUTIONS

Conceptualization, Y.G. and A.R.; Methodology, L.S., A.R. and Y.G.; Software, L.S. and A.R.; Validation, L.S.; Formal Analysis, L.S. and M.S.A.; Resources, A.R. and A.S.; Investigation, B.E., E.M.S. and Y.G.; Data Curation, L.S. and Y.G.; Writing - Original Draft, Y.G.; Writing - Review & Editing, A.R., L.S., Y.G., C.M., E.M.S., B.E. and M.S.A.; Visualization, L.S. and Y.G.; Supervision, Y.G., A.R. and C.M.; Project Administration, Y.G. and A.R.; Funding Acquisition, A.R., A.S. and C.M.

## METHODS

### Network architectures

In our framework, the nodes in the network represent genes, where the expression of a gene is regulated by the expression of other genes. Gene regulation is represented by directed edges in the network, e.g. if the expression of gene Y is regulated by the expression of gene X, then the network contains an edge from node X to node Y. These networks can be defined by adjacency matrices given by:

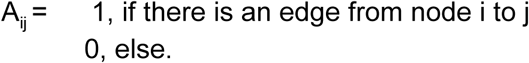

Any node in a network of size N can be connected with up to N-1 other nodes and in the case of self-loops, to N other nodes. Hence, the adjacency matrix A is of size N*N. This means that there are 2^N×N^ possible adjacency matrices for a network of size N - each of the possible N*N matrix entries can take on one of the values of 0 (no edge) and 1 (edge). For example a network of size 3 has 2^(3*3)^ = 512 possible network architectures.

For our analysis, we focus on symmetric networks, where we assume a relational identity between all nodes in a network. This implies the absence of a hierarchical structure within the mechanistic driver genes, and that all driver genes act equally in this respect (**Figure S1E**). The structural embedding of a node in its network could increase or decrease its ability of being involved in coordinated overexpression. To ensure for a relational identity, we define a set of symmetric networks, where the number of in- and outgoing edges within a node and across nodes is identical and either all nodes in a network have a self-loop or not. This leads to adjacency matrices of which the rows are cyclic permutations (to the right) with offset one of each other. For this, we first compute all possible vectors {0,1}^N^, in total 2^N^ vectors. From each of these resulting vectors, we create an NxN matrix by using the given (row) vector as template, and creating the other N-1 rows by cycling the prior row vector to the right by one step, where the right-most entry in the row vector is added to the (so far empty) left-most entry. By applying this permutation N-1 times, all possible cyclic permutations are captured within a matrix, and each node in the given network is completely relational identical. We make use of the *circshift* function in MATLAB to receive the possible cyclic permutations of the initial row vectors.

We further constrain the analysis to weakly-connected networks -- any node in a network has to be connected to at least one other node, without taking into account the directionality of the edges. In terms of the adjacency matrix, this implies:

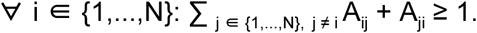

This restriction allows us to exclude the consideration of compositions of smaller and unconnected networks, which could otherwise lead to double counting. These subnetworks of smaller sizes are analyzed in the sets of networks of respective node sizes. To perform this operation, we analyze all the previously constructed adjacency matrices using the MATLAB function *conncomp(X,’Type’,’weak’)*, which assigns each node with a bin number according to the connected component of its underlying undirected graph. If all nodes of a network belong to the same bin number i.e. to the same connected component, the adjacency matrix encodes for a weakly-connected graph. Finally, we further restrict our analysis to non-isomorphic networks. Two networks are called isomorphic, if there exists a bijection from the edge space of one network to the other, such that any edge of one network is projected to a particular edge in the other network. In our case, the labeling of the nodes (gene 1, gene 2, …) in the networks is arbitrary and hence relabeling of nodes in an adequate fashion leads to identical networks in our analysis. To ensure that all the final networks analyzed are of a non-isomorphic set of networks, we test all networks with MATLAB’s function *isisomorphic*. We initiate the final set of networks with one adjacency matrix, and then sequentially test all other networks for isomorphism. If the given network is non-isomorphic to the current final set, it is added to the final set. Conversely, if the network is isomorphic to one of the networks in the final set, it is discarded.

By reducing the possible set to weakly-connected, non-isomorphic and symmetric networks, we greatly reduce the possible number of network architectures. For example, in the previous example, we had 512 possible network architectures for 3 nodes. By applying all the mentioned constraints (weakly-connected, non-isomorphic and symmetric), 4 network architectures remain (**Figure S1E**). We perform our analysis on networks of sizes 2, 3, 5 and 8 each consisting of 2, 4,10 and 80 network architectures, respectively, adding up to a total of 96 network architectures (**Figure S9**). In principle, our model can easily be extended to larger network sizes without the loss of generality.

### Model 2 - Transcriptional bursting-based stochastic network model

Our model is an expansion of the telegraph model, where DNA can take on one of the two states, active and inactive, e.g. based on the presence or absence of transcription factors. The active and inactive state directly translates into high and low rates of production of gene products, respectively. We add interaction terms to this model, where the expression of a gene influences the rate of DNA activation of another gene depending on how they are organized in a respective network. Here we use the number of mRNA as a faithful proxy for the number of proteins. In other words, we only model the number of mRNA counts and assume that any mRNA is immediately translated into one single functional protein after its translation. Therefore, the mRNA count determines the strength of the regulation. Here, we model the regulation of one gene by another using the Hill function, given by:

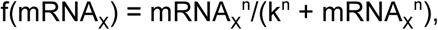

where mRNA_X_ is the mRNA count of gene X, n is the Hill coefficient and k is the dissociation constant, n,k > 0. The Hill coefficient determines the steepness of the Hill function, i.e., the extremeness of its switch-like effect. The dissociation constant determines the half-maximal value, f(mRNA_X_) = 0.5.

The reversible transitions between the inactive and active states, as well as the mRNA synthesis and degradation, are modeled by chemical reactions. For each gene, we have three chemical species - the DNA inactive state, the DNA active state and mRNA. These three species interact with one another according to the following 5 chemical reactions:

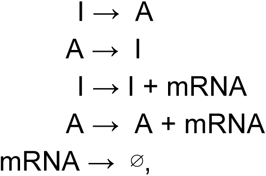

defining the corresponding stoichiometric matrix:

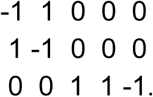

The stoichiometric matrix encodes the net change in each chemical species resulting from any of the chemical reactions where the chemical reactions are assumed to occur stochastically. Under the assumptions of the law of mass action, the probability of a specific molecular collision to occur in the infinitesimal time interval [t, t + dt) is proportional to the product of the molecule counts of the educt chemical species. The reaction propensity a_j_(x) for a given chemical reaction R_j_ and state x, determines the probability density function such that a_j_(x)dt gives the probability of the chemical reaction R_j_ taking place in dt, for small dt. Examples of reaction propensities for so called elementary reactions are given here:

| Reaction | Reaction propensity |
| --- | --- |
| $\emptyset \rightarrow \text{products}$ | $k$ |
| $X_i \rightarrow \text{products}$ | $kx_i$ |
| $X_i + X_j \rightarrow \text{products}$ | $kx_i x_j$ |

where k is called the reaction rate.

The gene regulation influences the reaction rate of the DNA activating chemical reaction. To explain the above-mentioned chemical reactions, we introduce eight rates/parameters:

| Parameter | Description |
| --- | --- |
| $r_{on}$ | The rate at which DNA is activated. |
| $r_{off}$ | The rate at which DNA is inactivated. |
| $r_{prod}$ | Synthesis rate of mRNA. |
| $r_{deg}$ | Degradation rate of mRNA. |
| $r_{add}$ | Parameter determining the contribution of the additional DNA activation rate upon gene regulation. |
| $d$ | Factor by which the mRNA synthesis rate is increased when in an active DNA state (in comparison to basal synthesis rate in DNA inactive state), $>1$. |
| $k$ | Dissociation constant of the Hill function. |
| $n$ | Hill coefficient. |

The full model description for one gene regulated by a single gene X is given below:

| Chemical reaction | Reaction rate | Reaction propensity |
| --- | --- | --- |
| $I \rightarrow A$ | $r_{on} + r_{add} * mRNA_X^n / (k^n + mRNA_X^n)$ | $(r_{on} + r_{add} * mRNA_X^n / (k^n + mRNA_X^n)) * I$ |
| $A \rightarrow I$ | $r_{off}$ | $r_{off} * A$ |
| $I \rightarrow I + mRNA$ | $r_{prod}$ | $r_{prod} * I$ |
| $A \rightarrow A + mRNA$ | $d * r_{prod}$ | $d * r_{prod} * A$ |
| $mRNA \rightarrow \emptyset$ | $r_{deg}$ | $r_{deg} * mRNA$ |

where I,A ∈ {0,1}, and I+A = 1, where I = 0 (A = 1) denotes that the DNA is in an active state and I = 1 (A = 0) denotes that the DNA is in an inactive state. mRNA_X_ is the mRNA count of gene X at the given time, r_on_ is the basal DNA activation rate, r_add_ is the additional activation rate due to gene regulation, r_off_ is the DNA inactivation rate, r_prod_ is the basal mRNA synthesis rate in the DNA inactive state, d denotes the increase in the mRNA synthesis rate when the DNA is in the active state, where d > 1, and r_deg_ is the mRNA degradation rate. The chemical reactions are identical for all N nodes in a given network of size N. The reaction rate of activation (I → A), composed of terms with parameters r_on_ and r_add_, is the only node-specific rate. It depends on the underlying network architecture and has to be adapted accordingly for each node, where the in-going edges of a node determine which gene regulations are active. This also corresponds to the adaptation of the standard telegraph model, highlighted in blue in the above rates. We model gene regulation additively: if there is more than one influencing gene, we add the Hill function terms of the respective genes. As an example, if the gene of interest is influenced not only by gene X, but by gene X and gene Y, the activation rate from above will expand to:

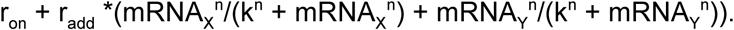

### Model 1 - Stochastic network model

Model 1 is a simple gene regulatory expression model, where mRNA can either be transcribed or degraded and the mRNA of a regulatory gene influences the transcription rate of a regulated gene. Here again, we assume the number of mRNA to be a faithful proxy for the protein number and hence, only model the mRNA expression of a gene. The gene regulation is modeled according to the Hill function (see Stochastic model of transcriptional bursting including gene regulation).

The synthesis and degradation are modeled by chemical reactions. For each gene, we have one chemical species, its mRNA, described by the following two chemical reactions:

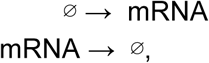

defining the corresponding stoichiometric matrix:

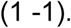

The full model description for one gene regulated by a single gene X is given below:

| Chemical reaction | Reaction rate | Reaction propensity |
| --- | --- | --- |
| $\emptyset \rightarrow \text{mRNA}$ | $r_{\text{prod}} + r_{\text{add}} * \text{mRNA}_X^n / (k^n + \text{mRNA}_X^n)$ | $r_{\text{prod}} + r_{\text{add}} * \text{mRNA}_X^n / (k^n + \text{mRNA}_X^n)$ |
| $\text{mRNA} \rightarrow \emptyset$ | $r_{\text{deg}}$ | $r_{\text{deg}} * \text{mRNA}$ |

where r_prod_ the basal mRNA synthesis rate, r_deg_ the mRNA degradation rate, r_add_ the additional synthesis rate due to gene regulation and mRNA_X_ the mRNA count of gene X at the given time.

The chemical reactions are identical for all N nodes in a given network of size N. The synthesis rate is a node-specific rate (see Stochastic model of transcriptional bursting including gene regulation). We model gene regulation additively (see Stochastic model of transcriptional bursting including gene regulation). For k we tested two different definitions: one closer and one further away from the low expression taking into account the intrinsic stochasticity. We therefore first run a test simulation with a random k for 1,000 time units and determine the standard deviation of the expression of the node denoted as ‘node 1’. K is latin hypercube sampled with the rest of the parameters with lower and upper boundary 100 and 1000. We set k to be:

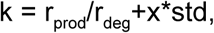

where std is the standard deviation of the expression of the node denoted as ‘node 1’ and x ∈ {3,5}. We then re-initiate the simulation with the adapted k value.

### Parameters

The goal of this framework is to model the emergence of rare transient coordinated high expression of several genes. The theoretical idea behind our model is that each time the DNA is in an active state, corresponding to a transcriptional burst, the steady-state of the mRNA count is shifted from r_prod_/r_deg_ to d*r_prod_/r_deg_. Accordingly, the mRNA attempts to reach its new steady-state which results in a rapid increase in their counts. Depending on the length of the transcriptional burst, which is exponentially distributed with rate parameter r_off_, the mRNA count is able to reach the new steady-state. We use this dynamical system behavior when modeling the rare coordinated overexpression. In principle, for most transcriptional bursts, the sudden mRNA increase should not initiate a DNA activation of its regulated genes; only in some rare cases, the transcriptional burst in one gene is long enough such that its mRNA count exceeds a certain threshold that may be able to affect the state of another gene locus on DNA. In turn, this can lead to an increased probability of the DNA states of its regulated genes to be activated and hence to an increased mRNA synthesis in the respective genes. This may lead to positive feedback loops within the network resulting in the transient coordinated overexpression of genes.

The threshold to be overcome by the mRNA count of a gene to make its gene regulation effective is given by the dissociation constant of the Hill function, k. k determines the ‘switching point’ from (almost) no gene regulation to (almost) complete gene regulation. Therefore, we define k to be a function of r_prod_, r_deg_ and d as follows:

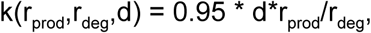

where d*r_prod_/r_deg_ gives the steady-state mRNA count of the respective regulating gene in the DNA active state. Here, we arbitrarily determine the threshold k to 0.95 of its high-expression steady-state to restrict the emergence of coordinated overexpression to being rare and for the system to demonstrate a significant difference between the low and high gene expression state. The simulations and the analysis are all performed according to this definition of k. We tested the robustness of this definition for a particular network 5.3 (**Figure S9**) where we performed the same simulations (for 100 latin hypercube sampled parameter sets (Supplementary Information ParSetsAnalysis.xlsx)) as for the final analysis as before using five different definitions of k:

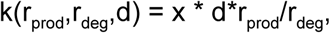

where x ∈ {0.75, 0.8, 0.85, 0.9, 1} (Supplementary Information ParSetsAnalysis.xlsx). This analysis shows that for x = 0.75, none of the 100 simulations show rare coordinated gene expression: the threshold leading to an effective gene regulation is exceeded too often: the regulated DNA states are activated, the high gene expression state emerges and we lose the rareness of the coordinated high gene expression event. The number of simulations showing rare behavior increases with increasing x, reaching its maximum for x = 0.95 (standard, 7 out of the 100 simulations show rare behavior). For x = 1 (high expression steady-state), we also see rare behavior in 7 out of 100 simulations, showing overlapping results in 6 out of the 7 simulations.

Together, we are left with a set of seven parameters consisting of: r_on_, r_add_, n, r_off_, r_prod_, d, r_deg_, which may be split into inter-gene (r_on_, r_off_, r_prod_, d, r_deg_) and intra-gene (r_add_, n) parameters and the dependent parameter k. Potentially, these parameter sets are node-dependent resulting in a N * 7-dimensional parameter space for a network of size N.

To emphasize the equality between the nodes, we use the same 7-dimensional parameter set for all nodes in a network (**Figure S1E**). Hence, the nodes are relationally and parametrically identical. This also allows us to directly compare the simulations of different network sizes, otherwise not possible, and to determine the effects of network size and architecture on the ability of forming the rare coordinated high-expression state. Therefore, we latin-hypercube sample 1000 parameter sets out of the parameter space with upper and lower boundaries (chosen arbitrarily, but typically spanning two orders of magnitude):

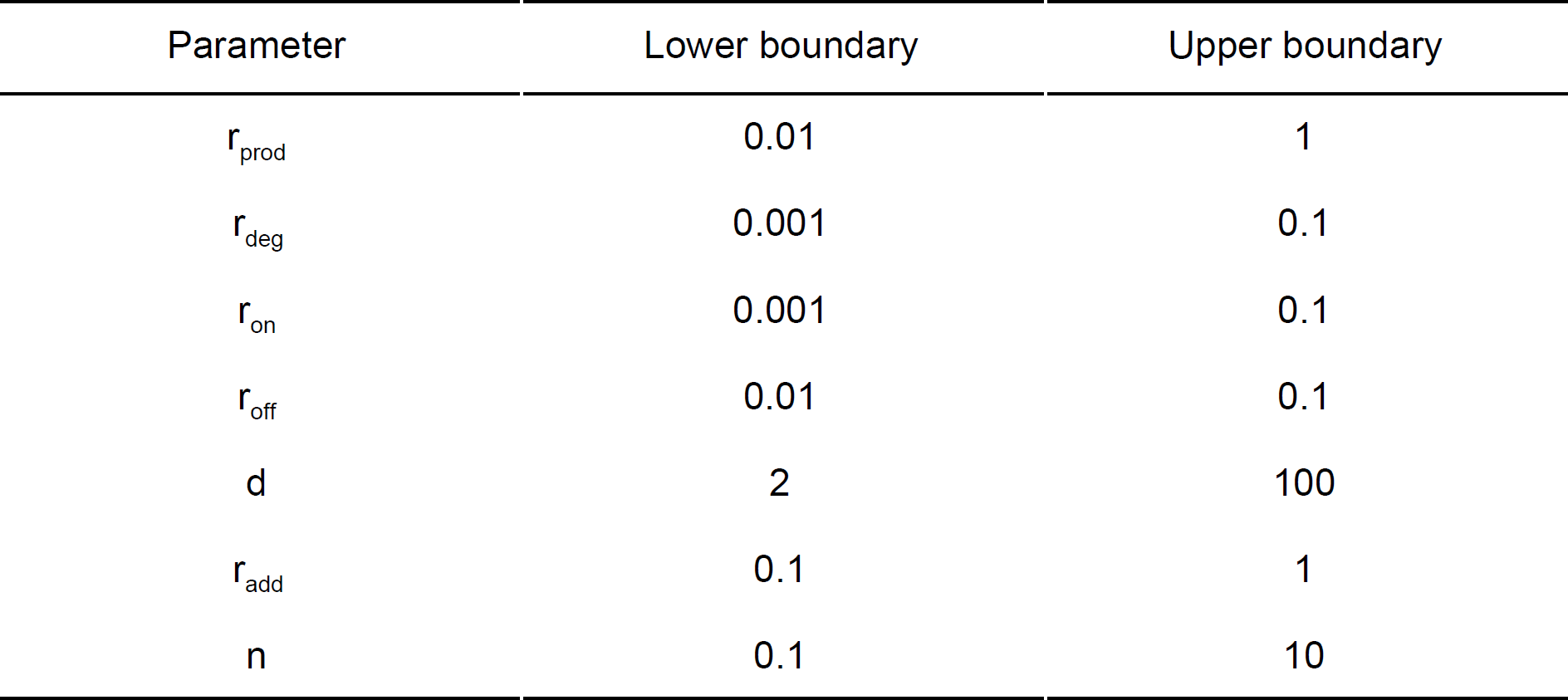

| Parameter | Lower boundary | Upper boundary |
| --- | --- | --- |
| $r_{\text{prod}}$ | 0.01 | 1 |
| $r_{\text{deg}}$ | 0.001 | 0.1 |
| $r_{\text{on}}$ | 0.001 | 0.1 |
| $r_{\text{off}}$ | 0.01 | 0.1 |
| $d$ | 2 | 100 |
| $r_{\text{add}}$ | 0.1 | 1 |
| $n$ | 0.1 | 10 |

by using the MATLAB function *lhsdesign_modified (*Khaled, N. Latin Hypercube (https://de.mathworks.com/matlabcentral/fileexchange/45793-latin-hypercube), MATLAB Central File Exchange. Retrieved May 5, 2018.). The 1000 parameter sets are shown in the Supplementary Information (ParSetsAnalysis.xlsx). For some plots, we used a y-axis break function in MATLAB(Mike,C.F.Break Y Axis (https://www.mathworks.com/matlabcentral/fileexchange/45760-break-y-axis), MATLAB Central File Exchange. Retrieved December 21, 2018.)

### Simulations

We simulated model 2 for a total of 96 network architecture (for all weakly-connected, non-isomorphic, symmetric networks of sizes 2, 3, 5 and 8 with 2, 4, 10 and 80 network architectures, respectively)(**Figure S9**), each for 1,000 sampled parameter sets. This results in a total of 96,000 simulations across four different network sizes. The simulations were performed according to Gillespie’s next reaction method and were computed for 1,000,000 time units, which is critical for capturing rare behaviors. For all simulations, the DNA state was initiated (t = 0) to be in its inactive state and the mRNA count was set to 20 for all nodes. This was determined arbitrarily. The mRNA counts quickly reach their low-expression steady state, such that we are certain that our analysis is not impaired by the given initial conditions. The simulations were implemented in MATLAB R2017a and R2018a. One single simulation of 1,000,000 time units took between 20 minutes and 9 hours depending on the parameter set and the network architecture. The complete simulations took over 1.5 months to run, where we parallelised all 96 networks and and let each of them run on four cores simultaneously.

### Simulation classes

We analyzed all of the 96,000 simulations, and assign them to the following four classes, initially by visual inspection, and subsequently by defined criteria (see below):

I - stably low gene expression

II - stably high gene expression

III - uncoordinated transient high gene expression

IV- rare, transient coordinated high gene expression

Therefore we constructed three criteria, for which all the simulations were tested. We primarily focus on the rare, transient coordinated high gene expression states, as defined by the following criteria:

#### 1) Coordinated high-gene expression state

We call a simulation to show coordinated high expression, if at least once within the 1,000,000 time unit simulation more than half of the mRNA counts are above a specified threshold (e.g. for 5 nodes, at least once three or more mRNA counts have to be above a defined threshold; for 8 nodes, at least once 5 or more mRNA counts have to be above a defined threshold). Similar to the definition of the dissociation constant k, we set the threshold to

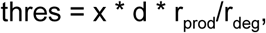

where d * r_prod_/r_deg_ gives the high-expression steady state. Again, we want to detect the rare occurrence of a large mRNA count deviation from the low-steady state and hence, set the threshold arbitrarily to 0.8 (see below for details on the choice of this value).

To compare our simulated results with the experimental data from a melanoma cell population, we split the 1,000,000 time unit simulations into 1,000 time unit sub-simulations, each accounting for a cell. Hence, we receive simulations of 1,000 cells for 1,000 time units. This procedure is justified by the ergodic theory. To show that sub-simulations of 1,000 time units are uncorrelated, we determine the autocorrelations for all 1,000 parameter sets of network architecture 3.2 (**Figure S9**) for up to 1,000 lags with the function *acf* (Price, C. (2011).Autocorrelation function(ACF) (https://www.mathworks.com/matlabcentral/fileexchange/30540-autocorrelation-function-acf), MATLAB Central File Exchange. Retrieved June 13, 2019.). For each of these, we determine the first lag at which the autocorrelation is below the upper 95% confidence bound. For 88.2% of all simulations, the first lag below the upper 95% confidence bound occurs before 1,000 lags.

#### 2) Rareness/transience

To mimic the results given by RNA-FISH in a melanoma population, where we only see a snapshot of the mRNA counts within a melanoma cell, we randomly determine a time point t_rand_, where t_rand_ ∈ [0,999] (uniformly distributed), at which we count the number of mRNA counts above the threshold (for each simulation t varies). We summarize the result of all 1,000 cells in a histogram, for which we expect a decrease with increasing mRNA count above the threshold.

#### 3) Heavy-tailed gene expression distributions

At the population level, the single mRNA distributions of marker genes show heavy-tails. We use the same time point t as sampled for criterion 2) and consider the mRNA counts of all genes. If we plot these in gene-dependent histograms, we expect to find right-skewed and unimodal distributions. Here, we use the MATLAB function *skewness(X)* for evaluating the right-skewness of the histogram, where skewness(X) > 0, denotes that the data is spread out more to the right of the mean. Skewness is defined as

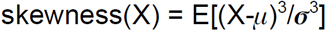

where *μ* is the mean of X, *σ* is the standard deviation of X and E(.) the expectation. For determining unimodality, we test whether the maximum of the last quarter of histogram bins with bin width of one is less than the minimum of the first quarter of histogram bins. Although this definition only characterizes a heavy-tailed distribution, we find it to be sufficient for this analysis.

Classes I and III, are both defined by criterion 1 only, where criterion 1 is not met in both cases. For class I, none of the genes in a network ever express above the given threshold. For class III, genes express above the given threshold but not once are more than half of the genes above the given threshold at any given time of the simulation. Only if a simulation is able to fulfill all three criteria, will we call it a simulation of class IV - rare transient coordinated high gene expression. If a simulation fulfills criteria 1, but fails to meet both other criteria, we classify it into class II.

To receive numbers of simulations in class IV - rare transient coordinated high expression - per network size, we randomly determine three different t_rand_, where each t_rand_ ∈ [0,999] (uniformly distributed) and evaluate all 96000 simulations for being in class IV at the respective snapshot (**Figure 2A**). Note that all these requirements are tested automatically using a script without manual/human intervention.

To show that criterion 3) is sufficient for defining heavy-tailed simulations in class IV in our analysis, we constrain criterion 3) further aiming to identify sub-exponentially decaying, heavy-tailed distributions more directly. We therefore reevaluate all simulations so far identified as class IV and compare their 99^th^ percentiles of their gene expression distributions with those of fitted exponential distributions (**Figure S2D**). We expect most of the 99th percentile of the gene expression distributions to be larger than the 99th percentile of the fitted exponentials. Due to the symmetry of the networks and the resulting similarity between the gene expression distributions (**Figure S2C**), we only consider node one in this analysis, without the loss of generality. To avoid that the fitted exponentials account for the heavy-tails, we constrain the fits to have a maximal bin number (bin size of one) within ∓ 1 of the maximal bin number (bin size one) of the expression distributions. We do so by sequentially increasing/decreasing the exponential parameter μ by steps of 10, sampling 1000 times from the resulting exponential distribution with the MATLAB function *exprnd*(μ,1,1000) and comparing the maximal bin number of the resulting histograms. We repeat this until the maximal bin number of the exponential distribution is within the predefined range of ∓ 1. As gene expression distributions with a large maximum bin are more similar to lognormal distributions with small variances and less to exponentials, we restrict this analysis to gene expression distributions with a maximum bin of ≤ 15 (**Figure S2E**). The threshold of a maximum bin of 15 was determined by considering the simulations and their exponential fits. We additionally discard simulations for which the optimization takes more than 1000 iterations or is producing non-positive parameter values.

Most (82%) of the 99th percentile of the gene expression distributions are above the diagonal, hence larger than the 99th percentile of the fitted exponential distributions (**Figure S2D**). With this we conclude, that criterion 3) sufficiently selects for sub-exponentially decaying heavy-tailed distributions.

We additionally, perform parts of the analysis again on two different levels of stricter stringency for criterion of heavy-tailed distributions (**Figure S3**):

A. All simulations fulfilling criteria 1) - 3) which additionally comply to the above mentioned analysis (maximum bin ≤ 15, 99th percentile of gene expression distribution > 99th percentile of fitted exponential, <1000 iterations to reach a ∓ 1 of the maximal bin number (bin size one) in the optimization for determining the exponential fit and producing non-positive parameter values) (**Figure S3E-H**).
B. All simulations fulfilling criteria 1) - 3) which additionally comply to the above mentioned analysis or have a maximum bin > 15 (**Figure S3A-D**).

The results are qualitatively very similar to the results we receive if we perform the analysis only on criteria 1) - 3) (**Figure S3**). The 6 and 7 rare coordinated high parameter sets identified by the more stringent analyses A) and B), respectively, are subsets of the original eight rare coordinated high parameter sets (**Figue 2G, FigureS3C** and **S3G**). Although the resulting optimized decision trees vary slightly, they still identify all three parameters, r_on_, r_add_ and r_off_, controlling rare transient coordinated states, as in the original analysis.

Together we conclude that the simple characterization of heavy-tailed distributions is sufficient for further analysis.

This analysis is a prerequisite for our further findings and statements. Due to its importance, we tested the robustness of this analysis with respect to the definition of the threshold, marking the mRNA count above which a gene is called to be in the high-gene expression state, and with respect to the number of mRNA counts required above the threshold to call it a coordinated high-gene expression state (both determining criterion 1). For the test network 5_3, we hence repeated the analysis for thresholds:

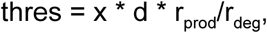

where x = 0.3 : 0.05 : 1 (here, for 100 latin hypercube sampled parameter sets (**Supplementary Information** ParSetsAnalysis.xlsx), and we only test for class IV). Decreasing the threshold down to 0.6 of the high-expression steady state does not change the set of simulations with rare behavior in comparison to the results for x = 0.8. Even a further decrease of the threshold (down to 0.3 of the high-expression steady state) manifests in a similar result: half of the simulations identified previously to show rare behavior are still classified as such. Hence, we keep x = 0.8 for the rest of this analysis (**Supplementary Information** ParSetsAnalysis.xlsx).

Next, for network 5.3 and the 100 parameter sets (**Supplementary Information** ParSetsAnalysis.xlsx), we repeated the analysis requiring at least 1, 2, 4, and 5 mRNA counts to be above the threshold at least once, in order for the simulation to fulfill criterion 1. The lower the required mRNA count, the more simulations fulfill criterion 1 (peaking at a required mRNA count of at least 1 with 11 out of the 100 simulations showing rare behavior according to this definition). This set of simulations entails the set of simulations fulfilling criterion 1 at our standard required mRNA count of at least 3 (7 out of 100 simulations). Hence, we keep our definition of coordinated overexpression to more than half the nodes being above the threshold.

### Network connectivity

We define a measure for the connectivity of the network architectures, where

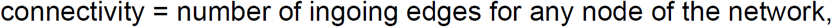

where a self-loop is also considered to be an ingoing edge. As we constrain our analysis to symmetric networks (same number of in-going edges for all nodes in a network per definition), we are able to define one single connectivity per network. This enables us to directly evaluate the impact of the connectivity of the network on the ability to form rare behavior.

### Quantitative Analysis

For each of the 96,000 simulations showing rare behavior we performed a quantitative analysis. First, we define a high expression region as a region which is initiated by the first mRNA count to exceed the threshold, terminated by the last mRNA count to drop below the threshold and requires to contain a coordinated high expression state (criterion 1: more than half the mRNA counts have to exceed the defined threshold) between the initiation and termination time points. Breaks of up to 50 time unit intervals are accepted due to the stochastic nature of the simulations. For example, in a 3 node network, where we require at least 2 mRNA counts to exceed the threshold for a coordinated high-expression state: the first mRNA count exceeds the threshold (initiation), then the second mRNA count exceeds the threshold (initiation of high-expression state) but then drops below the threshold for 50 time units before exceeding the threshold again, is still counted as one high-expression region. The length of 50 time units were defined arbitrarily. Due to the stochasticity of the system and the conservative definition of the threshold (located close to the high-expression steady state), we observe these temporary violations of criterion 1. In order, to create sensible statistics on the quantitative behavior of the simulations, this temporary relaxation of criterion 1 is necessary.

In the quantitative analysis we extract (i) the number of coordinated high-expression regions And (ii) the total time spent in a coordinated high-expression region (out of 1,000,000 time units) from all simulations showing rare behavior.

### Decision tree optimization, generalized linear models and constrained simulations

We classify all parameter sets into two classes, rare coordinated high parameter sets and non-rare coordinated high parameter sets, according to the percentage of total simulations per parameter set (96 simulations) in which rare behavior is observed. The threshold above which a parameter set is called a rare coordinated high parameter set is at 20%. More than 19 of the 96 simulations have to show rare behavior in order for a parameter set to be called a rare coordinated high parameter set. This threshold was set according to a summarizing histogram, in which we see a clear distinction between the two groups: the main body of the histogram being located below 20% and the few parameter sets deviating extremely from that main group (> 20%). According to this binary classification, we performed a decision tree optimization (MATLAB function fitctree).

To validate the results of the decision tree optimization, we used generalized linear models on all seven independent parameters r_on_, r_add_, n, r_off_, r_prod_, d and r_deg_ with the MATLAB function fitglm(X,Y,’Distribution’,’binomial’).

To validate that the parameter region determined by the decision tree optimization favors the formation of simulations with rare coordinated high states, we generate a new set of parameters constrained to values close to the minimal and maximal values of r_on_, r_add_ and r_off_ for the rare coordinated high parameter sets:

| Parameter | Lower boundary | Upper boundary |
| --- | --- | --- |
| $r_{prod}$ | 0.01 | 1 |
| $r_{deg}$ | 0.001 | 0.1 |
| $r_{on}$ | 0.001 | 0.025 |
| $r_{off}$ | 0.06 | 0.1 |
| $d$ | 2 | 100 |
| $r_{add}$ | 0.15 | 0.36 |
| $n$ | 0.1 | 10 |

where altered boundaries are indicated in blue. We latin hypercube sample 1000 parameter sets from that constrained parameter space. For all 1000 parameter sets we simulate 1000000 time units by Gillespie’s next reaction method for networks 3.2 and 5.3 (**Figure S9**). Each of these simulations was evaluated for having rare coordinated high states according to the three criteria (**Methods**, section Simulation classes).

### Entry and Exit mechanisms

#### Entering/Exiting of high expression region - Transcriptional bursts

For all of the simulations in class IV showing rare transient coordinated high gene expression - we analyze whether the durations of transcriptional bursts are coordinated with the entering and exiting of high expression regions (**Methods**, section Quantitative Analysis).

##### Entering high expression regions

For each of the defined high expression regions, we determine the entering gene - the gene corresponding to the gene count exceeding the threshold at the initial time point of the high expression region. We then extract all transcriptional bursts which do not start within a high expression region, determine their durations and classify them as either an entering burst or a non-entering burst. An entering burst is the last burst of a particular entering gene before or during its gene count exceeds the threshold. All other bursts are called non-entering bursts. We then perform a two-sample Kolmogorov-Smirnov test on the duration of the entering and non-entering bursts not in high expression regions with the MATLAB function *kstest2* at the significance level 0.05.

##### Exiting high expression regions

###### Transcriptional bursts

For each of the determined high gene expression regions we define an exiting region - the region between the first gene in the last quarter of the high expression region permanently leaving the high state (permanently having its gene count below the threshold for the rest of the high expression region) to the last time point of the high expression region. We again determine all transcriptional bursts - this time within the high expression regions. To exclude potentially prolonged entering bursts in this analysis, we only consider bursts which start within a high expression region. Also, for bursts exceeding the high expression region, we only account for their durations within the high expression region. If a burst overlaps with an exiting region for at least one time point we call the burst an exiting burst. All other bursts which are not overlapping with an exiting region are called non-exiting bursts. We apply the two-sample Kolmogorov-Smirnov test to the duration of the exiting and non-exiting bursts in high expression regions with the MATLAB function *kstest2* at the significance level 0.05.

#### Entering/Exiting of high expression region -Times

For all of the simulations showing rare transient coordinated high gene expression, we analyze the distributions of waiting times between genes entering and exiting the high expression region (see Quantitative Analysis).

##### Entering high expression regions

For all high expression regions, we determine the first time points at which the gene counts exceed the threshold (only for genes with a gene count exceeding the threshold during a particular high expression region at least once). We then consider the waiting times - the time interval between the ascending sorted time points of genes entering the high expression region. These distributions - at most N-1 distributions for a network of size N, one for each waiting time between the genes - are compared to exponential distributions by the Lilliefors test according to the MATLAB function *lillietest(X, ‘Distr’, ‘exp’)* at a significance level of 0.05.

##### Exiting high expression regions

For all high expression regions we determine the last time points at which the gene counts exceed the threshold (again, only for genes with a gene count exceeding the threshold during a particular high expression region at least once). We consider the waiting times and compare their distributions to exponential distributions by the Lilliefors test by applying the MATLAB function *lillietest(X, ‘Distr’, ‘exp’)* at a significance level of 0.05.

### Comparative Network Inference

Here we describe the computational techniques we used to infer the gene interaction network structure of the pre-drug and post-drug cells. When studying regulatory interactions between genes in a network, it can be useful to abstract the problem into a graph theory framework. Let us assume a set of N genes, with the expression level of each gene represented by the random variable *X*_*i*_, with *i* ∈ {*1,…,N*}. The network of interactions between genes can then be represented as a graph of *N* nodes. An edge *X*_*i*_ *→ X*_*j*_ signifies a regulatory relationship in which X_i_ either upregulates or downregulates *X*_*j*_ (Singh et al., *2012).*

The computational challenge of network inference is to uncover the true edges of the gene interaction network from statistical relationships between gene expression levels. Many different algorithms, often based on mutual information, conditional probability, or regression analysis, have been developed for this purpose (Huynh-Thu and Sanguinetti, 2019; Saint-Antoine and Singh, 2019; Singh et al., 2012). The output of an inference algorithm is a matrix of edge weights, which we will call W with dimensions NxN. In this matrix, the element *w*_*ij*_ is a measure of how confident we can be that the edge *X*_*i*_ *→ X*_*j*_ exists in the network. A final network prediction will typically set a threshold for edge weights, and exclude any edges that fall below the threshold. Edges *X*_*i*_ *→ X*_*i*_, called “self-edges”, are typically excluded for the final network prediction, except in cases when temporal data is being analyzed. Since we are using atemporal expression data in this analysis, self-edges will be excluded from our analysis.

It is common to judge a network inference algorithm’s reliability by testing it on a “gold standard” dataset, for which the true structure of the network is already known, to see how well it can recover the real edges from the expression data (Huynh-Thu and Sanguinetti, 2019). For this manuscript, we have chosen to use the Phixer algorithm (Singh et al., 2012), based on its impressive performance when benchmarked on the DREAM5 Challenge gold standard datasets (weblink: http://dreamchallenges.org/project/dream-5-network-inference-challenge/; last accessed: 05/06/2019).

#### Phixer

Phixer computes edge weights using the phi-mixing coefficient. For discrete random variables X and Y taking values in sets A and B, the phi-mixing coefficient Φ(X|Y) is defined as:

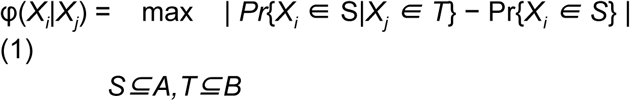

We then assign Φ(*X*_*i*_|*X*_*j*_) as the weight of the edge *X*_*j*_ *→ X*_*i*_. The phi-mixing coefficient is an asymmetric measure, so the weight of the edge *X*_*i*_ *→ X*_*j*_ may be different (Singh et al., 2012). The original Phixer algorithm includes a pruning step, which attempts to correct for false positives by minimizing redundancy in the network. For every possible triplet of nodes *X*_*i*_, *X*_*j*_, and *X*_*k*_, the following inequality is checked:

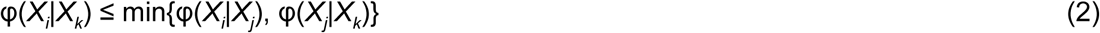

If Equation 2 holds, the edge *X*_*k*_ *→ X*_*i*_ is eliminated. However, previous work has found that the pruning step, though theoretically sensible, typically reduces accuracy in practice (Saint-Antoine and Singh, 2019), possibly due to the prevalence of redundant connections, such as feed forward loops, in gene regulatory networks. So, we removed this part of the algorithm in order to achieve the highest possible level of accuracy.

The Phixer software is available online at the creator’s Github page: https://github.com/nitinksingh/phixer/ (last accessed: 05/06/2019). We used the original C code, and kept the default parameter values the same, except for changing “NROW” to 19 and “TSAMPLE” to 4000, to reflect the dimensions of our input data files. The original Phixer code includes, by default, 10 bootstrapping runs, as well as a built-in procedure for binning the raw data, which we did not alter. We removed the pruning step from the code, but otherwise left the edge weight calculation process unchanged.

#### Data description

The two pre-drug datasets are referred to as NoDrug1 and NoDrug2 in the supplementary data files (**Supplementary Information** PhixerData.xlsx). The datasets containing clusters of resistant cells after four weeks of drug exposure are referred to as Fourweeks1-cluster1, Fourweeks1-cluster2, etc. where we differentiate between Fourweeks1 with four clusters and Fourweeks2 with three clusters. Details of how these datasets were acquired are presented in (Shaffer et al., 2017).

#### Bootstrapping controls

We found that the Phixer algorithm tends to predict more connections for larger sample sizes, even when the samples are taken from the same dataset. To correct for this issue and control for the differences in their original sample sizes, we bootstrapped the original datasets into 4000-sample datasets before performing the Phixer analysis. The number 4000 was chosen arbitrarily; bootstrapped sample sizes of 1000, 2000, and 6000 also appeared to produce similar results.

#### Randomized controls

For each size-controlled dataset to be analyzed, we created a randomized control consisting of permutations of each gene column from the original dataset (**Supplementary Information** PhixerData.xlsx). We then performed the Phixer analysis on these randomized controls. The resulting edge weight distributions give us a baseline or control edge weight for Phixer that, in principle, reflects potential false positives. We found that in the controls, nearly all of the predicted edge weights were below 0.45 (**Figure S7D**). Therefore, we decided to choose 0.45 as a threshold for our non-control analysis, thus eliminating edges that could have been predicted by chance alone.

Finally, since our analysis contains two stochastic elements (the bootstrapping to correct for the sample size issue and the bootstrapping step in the Phixer algorithm itself) we had to be sure that the observed differences in connectivity were not due to chance. For each dataset, we ran the entire analysis (including both the bootstrapping size correction and the Phixer algorithm) 1000 times, and provide the distributions of the number of edges with weight greater than 0.45 (**Supplementary Information** PhixerData.xlsx).

#### Asymmetric network architectures or parameter sets

To test the generality of our results, we generate asymmetric simulations. We introduce asymmetry in both network architectures and the parameter sets.

#### Asymmetric network architecture

We randomly determine a weakly-connected but asymmetric five-node network (**Figure S4A**). We simulate this network with 100 parameter sets which are latin hypercube sampled out of the same parameter space as the 1000 parameter sets of the main analysis. Out of these 100 simulations, two simulations are classified as showing rare, transient coordinated high gene expression (fulfills all three criteria in **Methods**, section Simulation classes, **Figure S4B** and **S4C**).

#### Asymmetric parameter sets

For the main analysis, we use the same parameter set, consisting of seven independent parameters (**Methods**, section Parameters), for all nodes in a network. We introduce asymmetry by assigning each node in a network a separate set of parameters. Hence, we latin-hypercube sample 100 parameter sets out of a 7 × N parameter space, where N is the number of nodes of the network, with the MATLAB function *lhsdesign_modified*. Due to the high dimensionality, we here confine the parameter space to:

| Parameter | Lower boundary | Upper boundary |
| --- | --- | --- |
| r_prod | 0.01 | 1 |
| r_deg | 0.001 | 0.1 |
| r_on | 0.001 | 0.1 |
| r_off | 0.001 | 0.1 |
| d | 2 | 100 |
| r_add | 0.2 | 0.4 |
| n | 5 | 10 |

where the changes in the boundaries are highlighted in blue. We confine the parameter space according to the clustering of rare coordinated high parameter sets. In total, six parameter sets give rise to rare-states more frequently than others for all 96 networks. Only two out of the seven independent parameters, r_add_ and n, show a strong correlation with the rare coordinated high state producing parameter sets as determined by a decision tree optimization. The boundaries in the table above are formed according to these decision tree boundaries in which five out of the six rare coordinated high state producing parameters lie (**Supplementary Information** ParSetsAnalysis.xlsx).

For these 100 parameter sets, we generated simulations for five-node network 5.3 (**Figure S4D**). Out of the resulting 100 simulations, we find two showing rare, transient coordinated high gene expression (fulfills all three criteria in **Methods**, section Simulation classes, **Figure S4E** and **S4F**).

## Supplementary Information

ParSetsAnalysis.xlsx

PhixerData.xlsx

