## Supplementary Information and Figures for "Gene networks with transcriptional bursting recapitulate rare transient coordinated expression states in cancer"

We decided to develop a network-based framework that models the cell-intrinsic biochemical interactions. One of the first goals we had was to identify the minimal set of biochemical reactions that constitutes this network model. We asked whether a simple network model lacking gene activation step (Model1), i.e. with constitutive mode of gene expression, is sufficient to capture rare coordinated high states (**Figure S1B**; **STAR Methods**, section Model 1)? Or that we need to incorporate gene activation step *via* transcriptional bursting (Model 2) at each node, a phenomenon in which genes flip reversibly between transcriptionally active and inactive state regulated by the binding of a transcription factor(s) (**Figure 1B**; **STAR Methods**, section Model 2)?

In terms of chemical reactions, the critical difference between the two models is that, while in Model 1 the gene is transcribed as a Poisson process with a single rate, $r_{prod}$ (**Figure S1B**), in Model 2, a gene can reversibly switch between active ($r_{on}$) and inactive state ($r_{off}$), where binding of the transcription factor at a gene locus defines the effective rate of gene production (**Figure 1B**). Specifically, when inactive, the gene is transcribed as a Poisson process at a basal rate ($r_{prod}$); when active, this rate becomes higher ($d \times r_{prod}$, where $d > 1$). For both the models, we modeled degradation of the gene product as a Poisson process with degradation rate $r_{deg}$. For both the models, the inter-node interaction parameter, $r_{add}$, has a Hill-function-based dependency on the gene product amount (Hill coefficient n) of the respective regulating node to account for the multistep nature of the interaction (**Figure 1B** and **S1B**). All chemical reactions, propensities, and model parameters are presented in **STAR Methods**. To test these two models, we used Gillespie's next reaction method (Gillespie 1977) and simulated test cases of small networks (of two or three nodes) for a range of parameters.

For a vast majority of the networks and parameter combinations, Model 1 either produced always low or always high expression states (**Figure S1C**). In some cases, while Model 1 could indeed produce a transition from low to high expression states, the transition happens for all gene products at the same time (**Figure S1C**). However, this model is not consistent with the experimental observations; in particular, if a cell is positive for one marker gene, then it is more likely to be positive for another marker gene, but not necessarily so (**Figure S1D**) (Shaffer et al. 2017). Furthermore, this mode of transition resulted in bimodal distributions of cellular state as determined by the amount of gene product (**Figure S1D**), which is different from the rare nature of the transitions, as reflected by the heavy-tailed distributions of gene products observed in melanoma. Model 2, which incorporates transcriptional bursting-dependent activation of a node (gene), also produced a range of gene expression states (**Figure 1C-1F**). Importantly, this model was able to faithfully capture the qualitative features of the experimental data i.e. rare, transient, and coordinated high expression states (**Figure 1F**). In contrast to Model 1, Model 2 captures another property of the experimental data, i.e. if one gene is in the high expression state, the other genes in the network are likely to be in high expression state, but not always (**Figure 2B** and **S2B**). Based on these initial observations, we decided to pursue Model 2 systematically and simulated networks of different sizes and architectures across a broad range of model parameters.

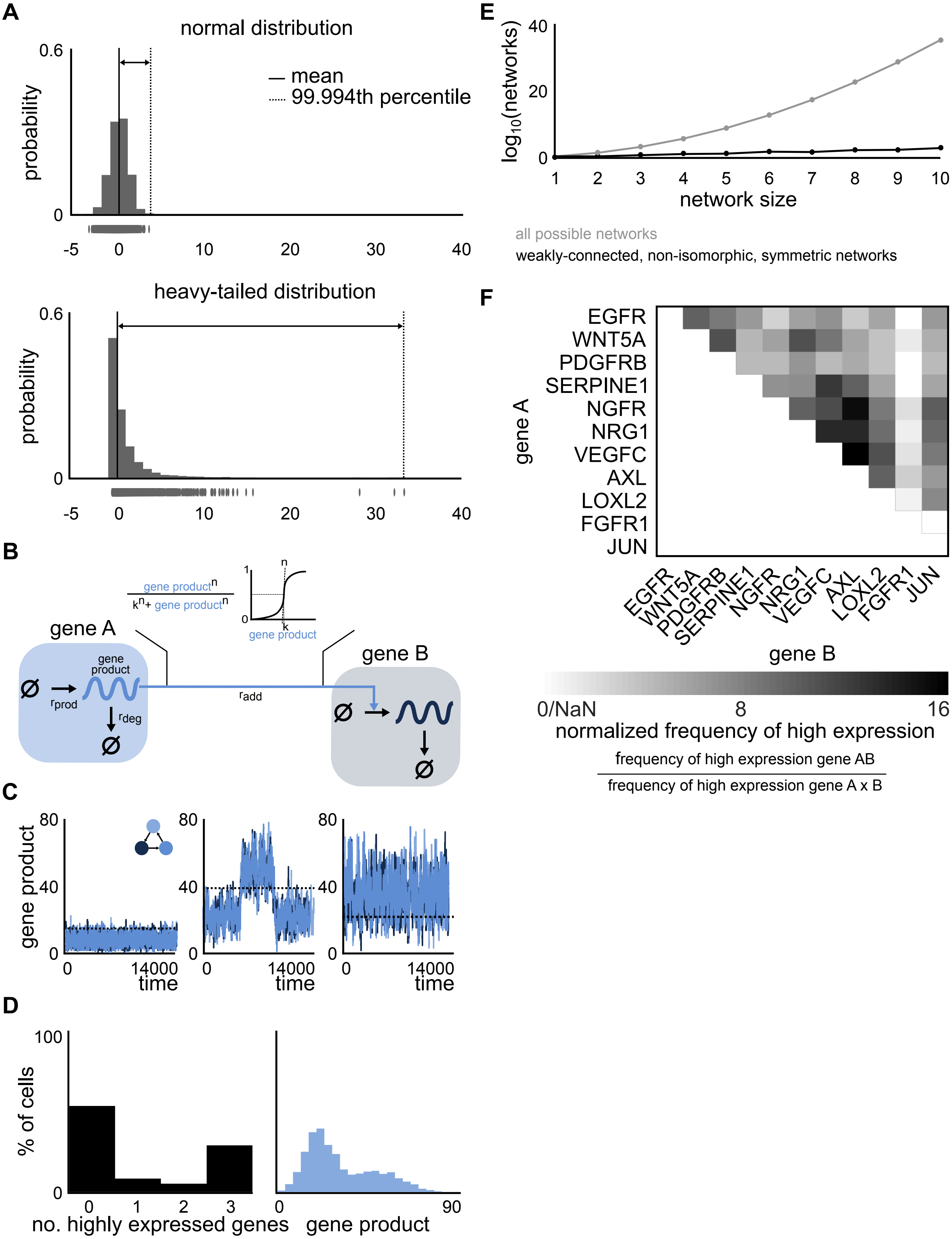
A
normal distribution
— mean
99.994th percentile
probability
heavy-tailed distribution
B
gene A
gene B
gene product
rprod
rdeg
radd
C
gene product
time
D
% of cells
no. highly expressed genes
gene product
E
log10(networks)
network size
all possible networks
weakly-connected, non-isomorphic, symmetric networks
F
EGFR
WNT5A
PDGFRB
SERPINE1
NGFR
NRG1
VEGFC
AXL
LOXL2
FGFR1
JUN
gene A
gene B
0/NaN
8
16
normalized frequency of high expression
frequency of high expression gene AB
frequency of high expression gene A x B

**Figure S1. Related to Figure 1 and STAR Methods.**
(A) A normal distribution with $\mu = 0$ and $\sigma^2 = 1$ (top) and a heavy-tailed distribution, here a log-normal distribution, with $\mu = 0$ and $\sigma^2 = 1$ (bottom). The 99.994th percentiles indicate that rare events are further from the bulk for heavy-tailed distributions than for normal distributions.
(B) Schematic of Model 1 - Stochastic network model. mRNA is either transcribed at rate $r_{prod}$ or degraded with rate $r_{deg}$. Gene regulation is modeled by a Hill function, where the gene expression of the regulating gene A increases the production rate of the regulated gene B.
(C) Depending on the network architecture and the parameters of the gene expression model, we observe either stably low expression (left), stably high expression (right) or transient coordinated high expression (middle).
(D) The distributions of simultaneously overexpressed genes and the gene products at the population level of (C middle) show bimodal distributions and are inconsistent with the observations in melanoma cells.
(E) The subset of weakly-connected, non-isomorphic, symmetric networks decreases the testable architecture space by many orders of magnitude.
(F) No clear driver gene or hierarchy is apparent from the two-dimensional RNA FISH experimental data.

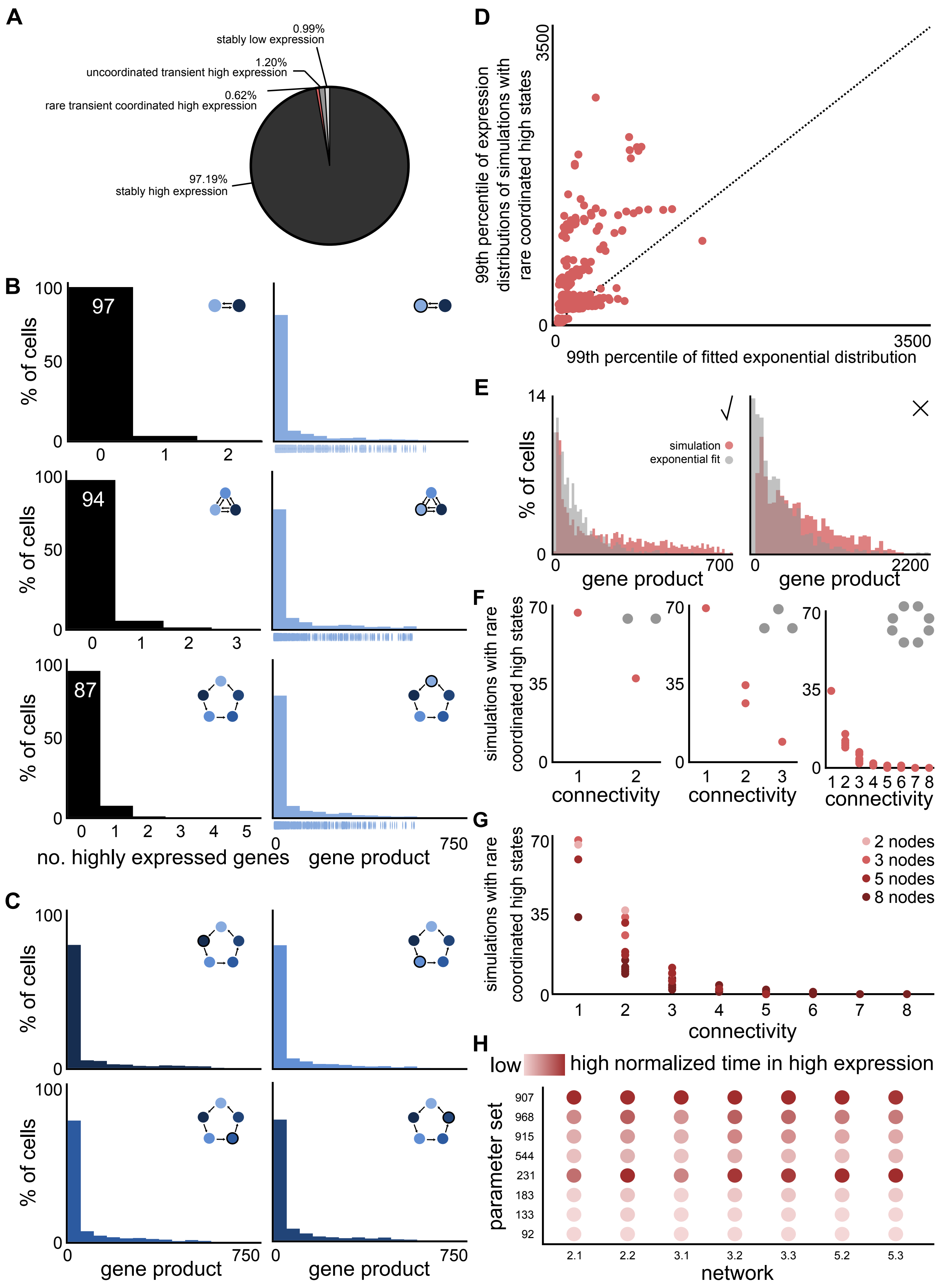

A
0.99%
stably low expression
1.20%
uncoordinated transient high expression
0.62%
rare transient coordinated high expression
97.19%
stably high expression
B
97
94
87
% of cells
no. highly expressed genes
gene product
C
gene product
D
99th percentile of expression distributions of simulations with rare coordinated high states
99th percentile of fitted exponential distribution
E
simulation
exponential fit
% of cells
gene product
F
simulations with rare coordinated high states
connectivity
G
2 nodes
3 nodes
5 nodes
8 nodes
connectivity
H
low
high normalized time in high expression
parameter set
network

**Figure S2. Related to Figure 2 and STAR Methods. Rare coordinated high states depend on network connectivity and model parameters but not on network size.**
(A) 0.62% of simulations show rare transient coordinated high expression.
(B) The simulated distributions of simultaneously highly expressed genes and expression are qualitatively similar to data from a pre-resistant melanoma population ubiquitously in networks with different numbers of nodes. Shown for a two node (top), three node (middle) and eight node network (bottom).
(C) The gene expression distributions of all five nodes (the gene expression distribution of node one is shown in (B)) are qualitatively similar.
(D) The tails of the simulated distributions for gene expressions are fatter than of fitted exponential distributions (see STAR Methods).
(E) Simulated gene expression distributions deviating too much from exponential distributions are discarded in the analysis shown in (D).
(F) Increasing connectivity within all networks of sizes two (left), three (middle) and eight (right) leads to a decrease in the number of simulations with rare coordinated high states.
(G) For connectivities of seven and eight, none of the networks showed simulations with rare coordinated high states.
(H) Simulations of particular parameter sets across different network architectures and sizes show similar (normalized) time in high expression relative to other parameter sets.

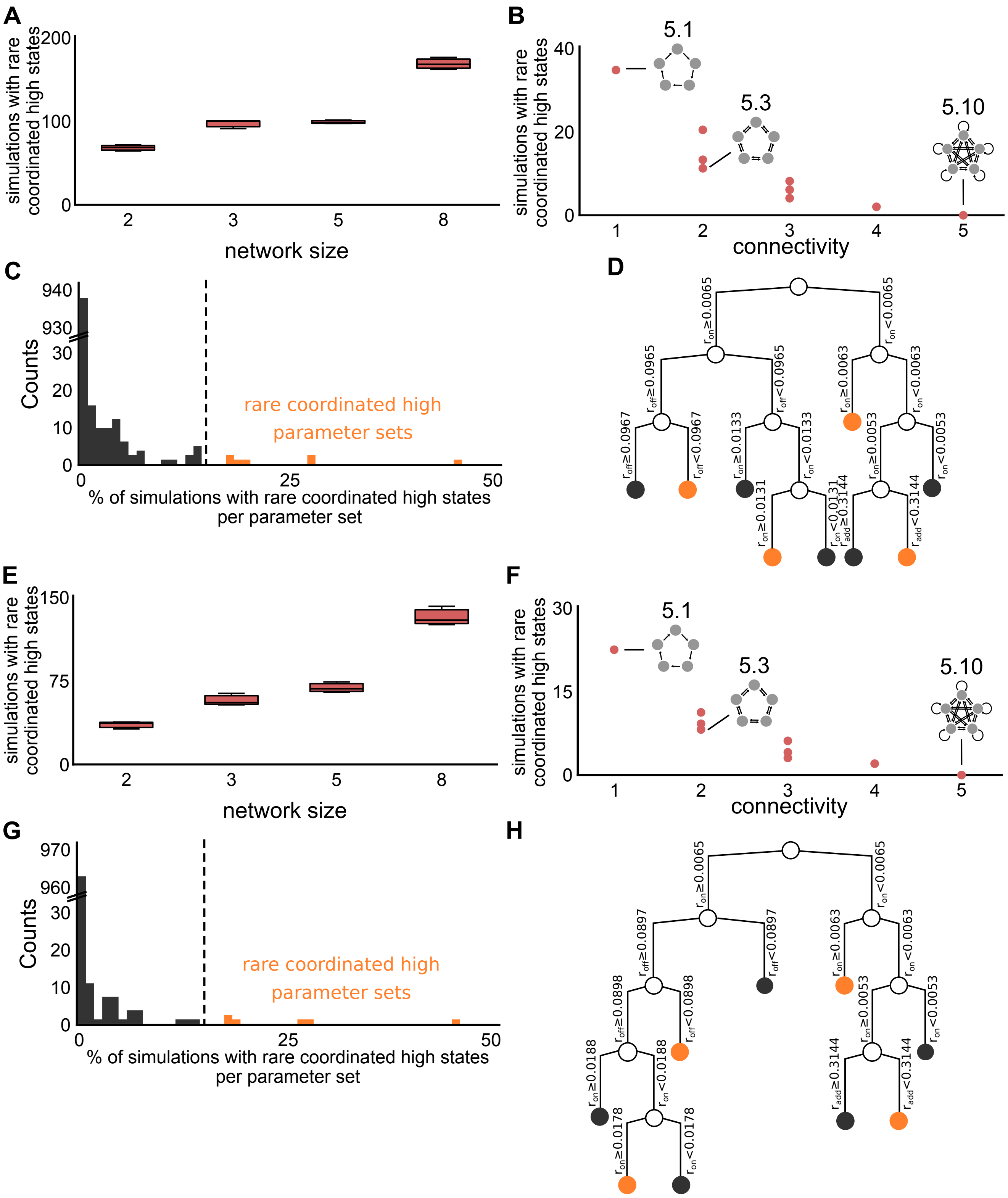


**Figure S3. Related to Figure 2, Figure S2 and STAR Methods.** Two levels of stringencies for the definition of heavy-tailed distributions show qualitatively similar results (A-D and E-H) to each other and to the stringency defined in main text (Figure 2).

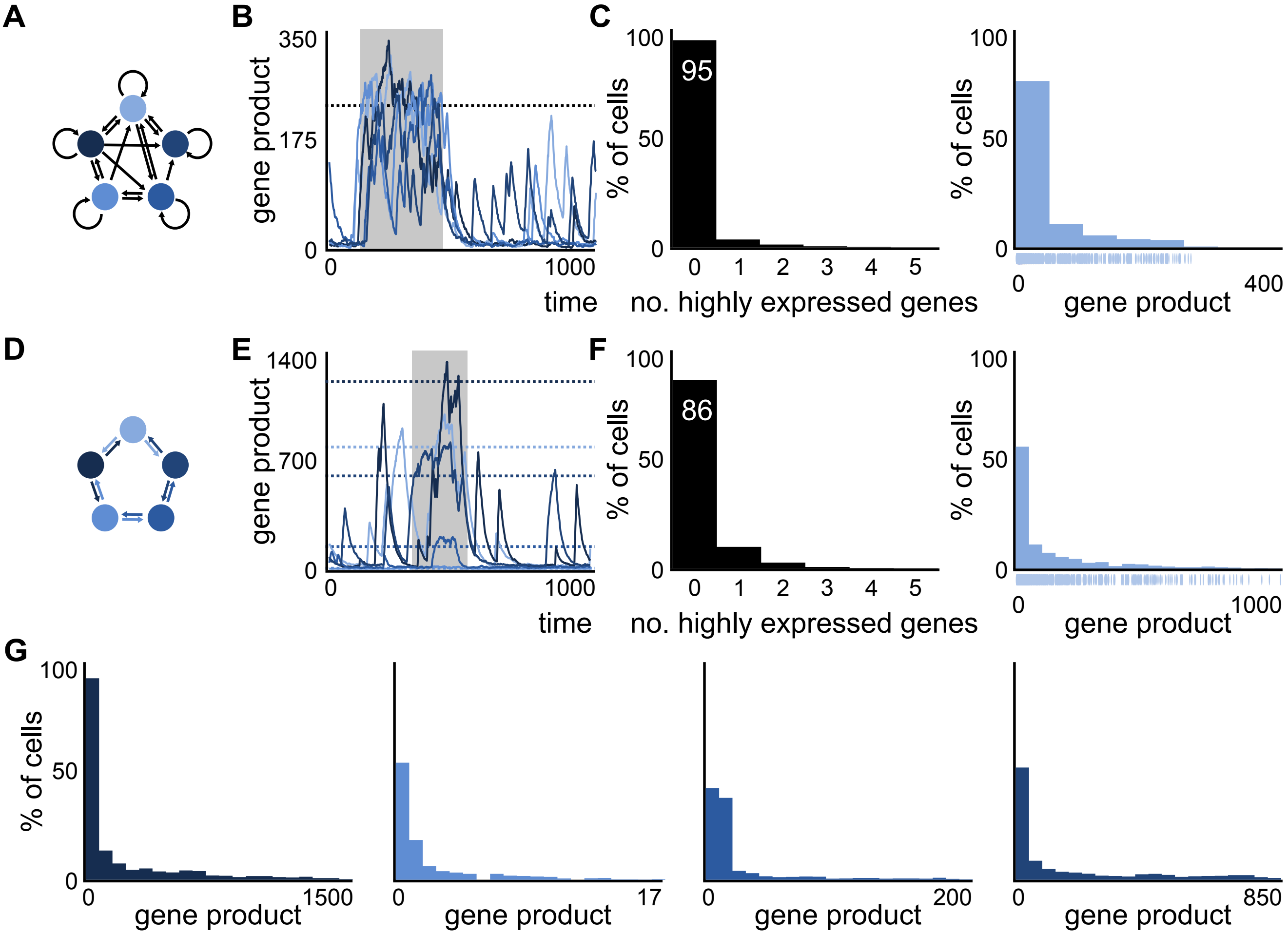


**Figure S4. Related to Figure 2 and STAR Methods. Simulations of asymmetric networks and asymmetric parameter sets show high coordinated states.**

(A-C) Asymmetric network architecture (A) with corresponding simulation (B) and distributions of simultaneously overexpressed genes and gene expression (C). The distributions show qualitatively the same behavior as drug naive melanoma cells.

(D-F) Symmetric network architecture and an asymmetric parameter set (D) with corresponding simulation (E) and distributions of simultaneously overexpressed genes and gene expression (F) . The distributions show qualitatively the same behavior as drug naive melanoma cells.

(G) The gene expression distributions of all five nodes (the gene expression distribution of node one is shown in (F)) generated with an asymmetric parameter set display different levels of heavy-tails.

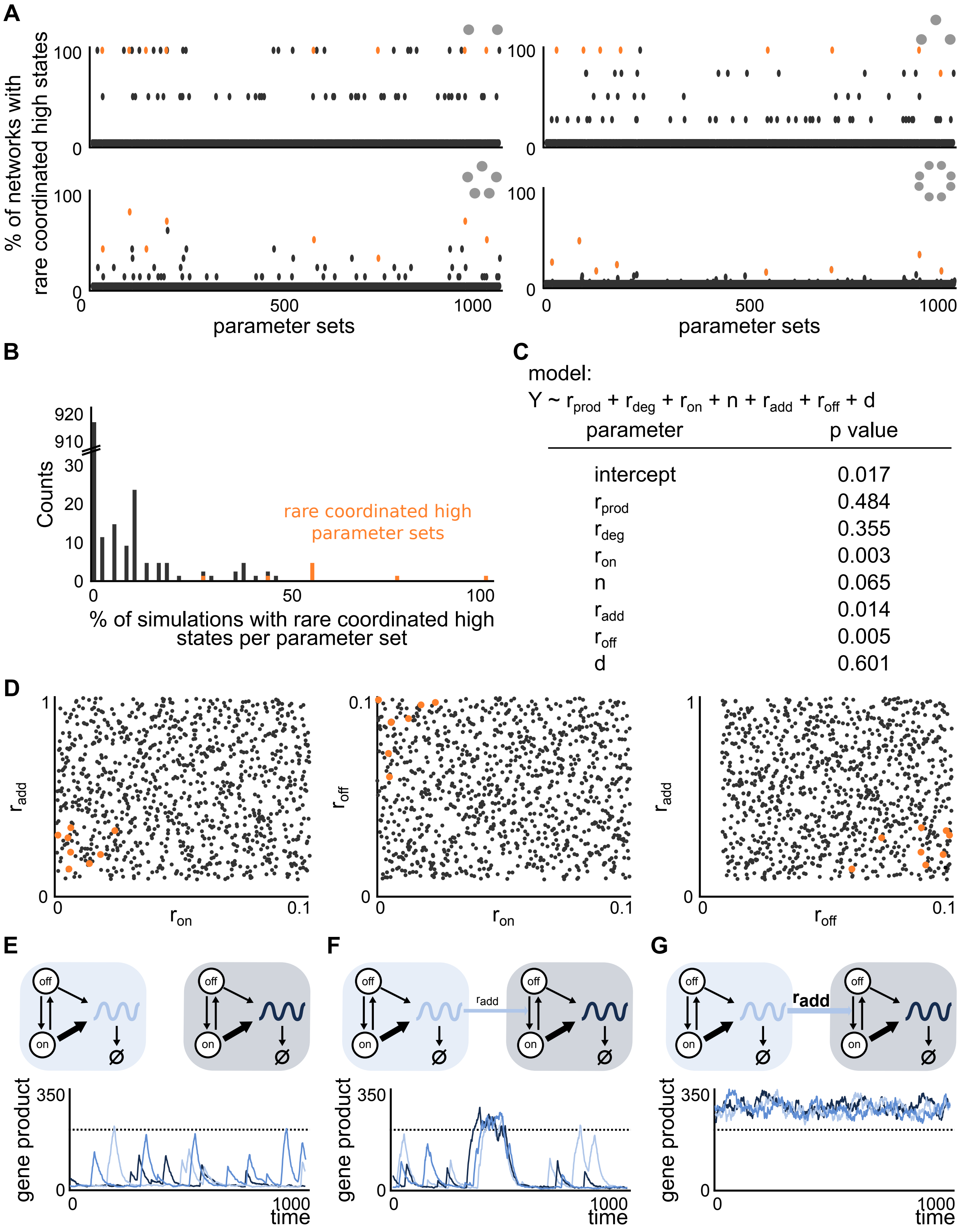
A
% of networks with rare coordinated high states
100
0
500
1000
parameter sets
B
Counts
920
910
30
20
10
0
0
50
100
rare coordinated high parameter sets
% of simulations with rare coordinated high states per parameter set
C
model:
Y ~ r_prod + r_deg + r_on + n + r_add + r_off + d
parameter | p value
intercept | 0.017
r_prod | 0.484
r_deg | 0.355
r_on | 0.003
n | 0.065
r_add | 0.014
r_off | 0.005
d | 0.601
D
r_add
r_on
r_off
1
0.1
0
E
F
G
off
on
r_add
gene product
350
0
1000
time

**Figure S5. Related to Figure 2. Three out of seven parameters give rise to the rare coordinated high states more frequently than others**.

(A) The rare coordinated high parameter sets (orange) give rise to rare coordinated high states more frequently than others in any given network of sizes two, three, five, and eight (from top left to bottom right).

(B) The rare coordinated high parameter sets are consistent even when confining the analysis to networks with connectivity of three or less.

(C) Analysis of the parameter sets by the generalized linear model where the model specification, parameters, and the respective p values are shown. Parameters with p value less than 0.05 are considered significant.

(D) Two dimensional representations of all tested 1000 parameter sets for $r_{on}$ - $r_{add}$ , $r_{on}$ - $r_{off}$ and $r_{off}$ - $r_{add}$ show that the rare coordinated parameters are narrowly constrained in the respective 2D spaces (orange).

(E-G) Increasing parameter $r_{add}$ leads to more stable high expression shown for $r_{add}$ = 0 (E), $r_{add}$ = 0.29 (F) and $r_{add}$ = 100,000 (G).

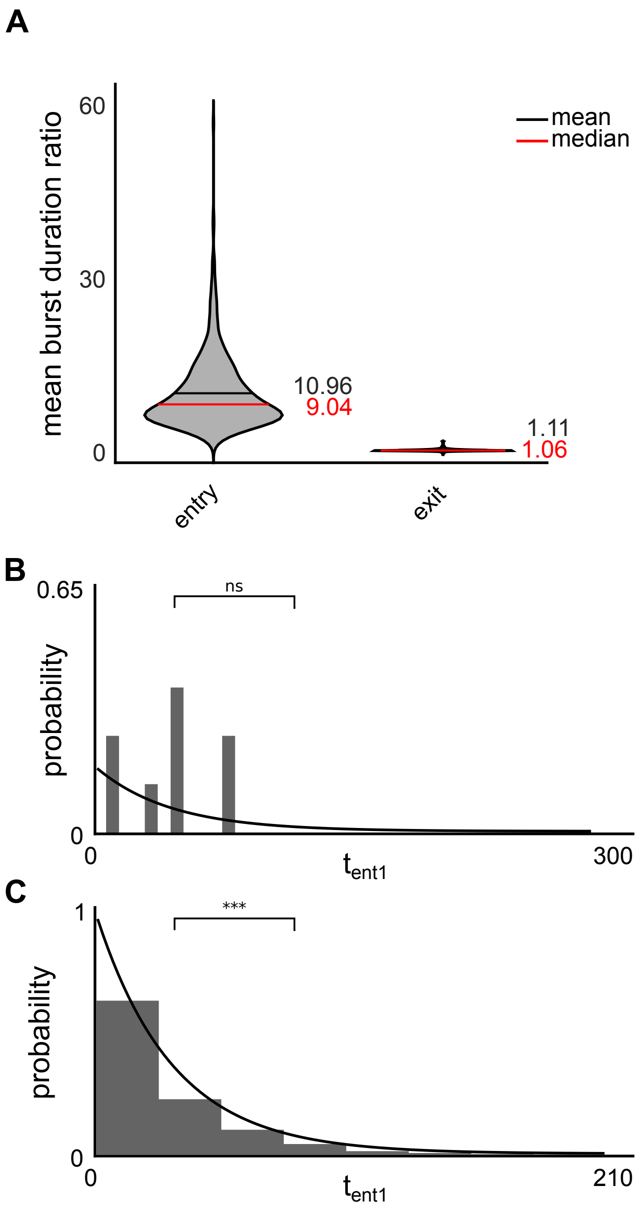


**Figure S6. Related to Figure 3.**

(A) (Left) Ratio of burst durations during entry time-point and baseline time-region. (Right) Ratio of burst durations during the high time-region and exit time-region. Ratio close to 1 suggests no difference between the two regions.

(B) Representative plot of distribution that satisfies the Lilliefors test corresponding to Figure 3D.

(C) Representative plot of distribution that rejects Lilliefors test corresponding to Figure 3E.

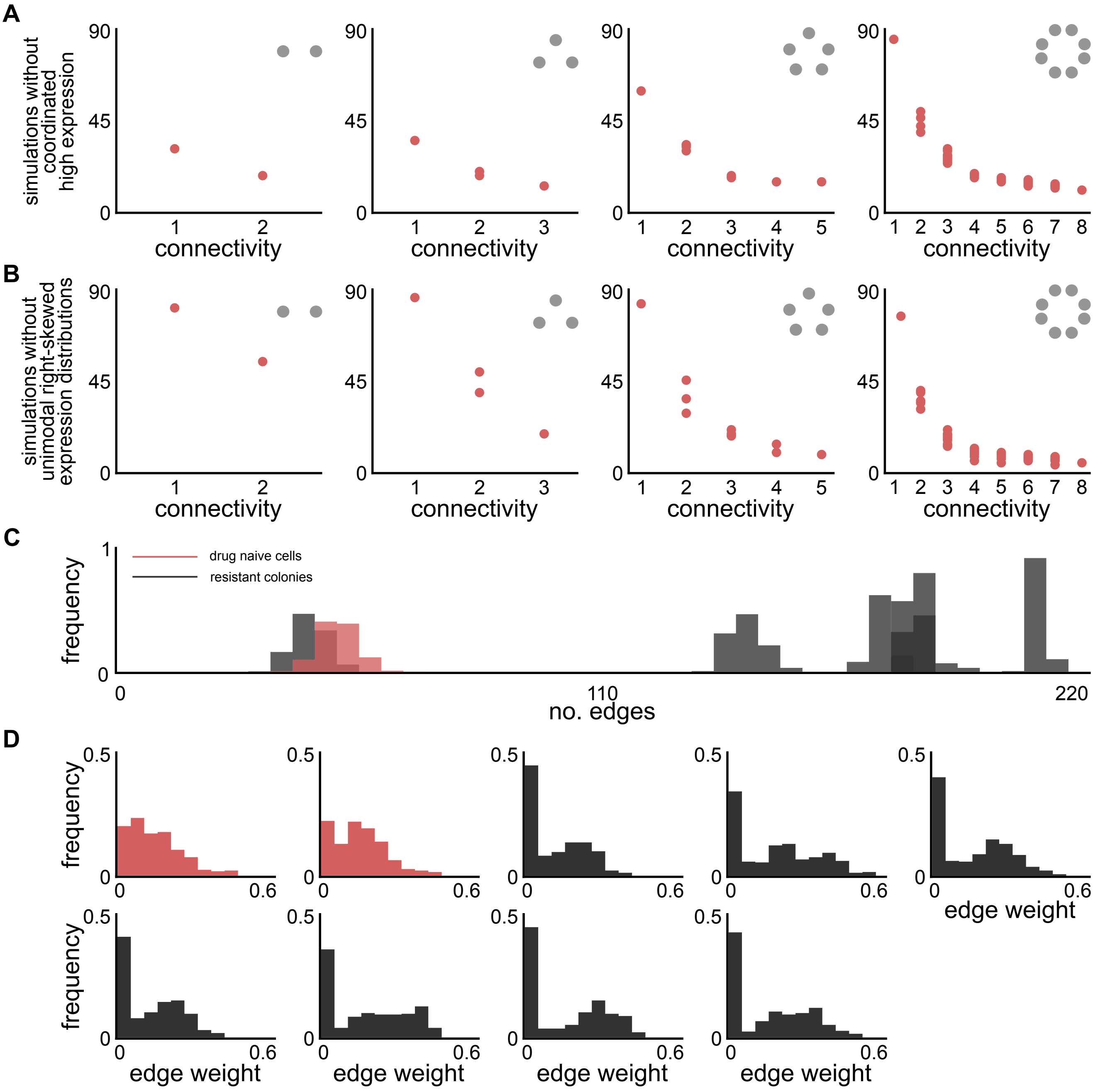


**Figure S7. Related to Figure 4 and STAR Methods. With increasing connectivity, simulations are more likely to enter the coordinated high state but are no longer able to leave it.**

(A) With increasing connectivity, less simulations fail to show coordinated high expression i.e. more simulations show coordinated high expression, shown for networks of size two, three, five and eight (left to right).

(B) With increasing connectivity, less simulations show heavy-tailed expression distributions i.e. more simulations show bimodal distributions and stable high expression, shown for networks of size two, three, five, and eight (left to right).

(C) The number of edges in the inferred gene regulatory networks are higher in 6/7 resistant colonies than in the two biological replicates of drug naive cells.

(D) For randomized controls, the edge weight is below 0.45, shown for all biological replicates.

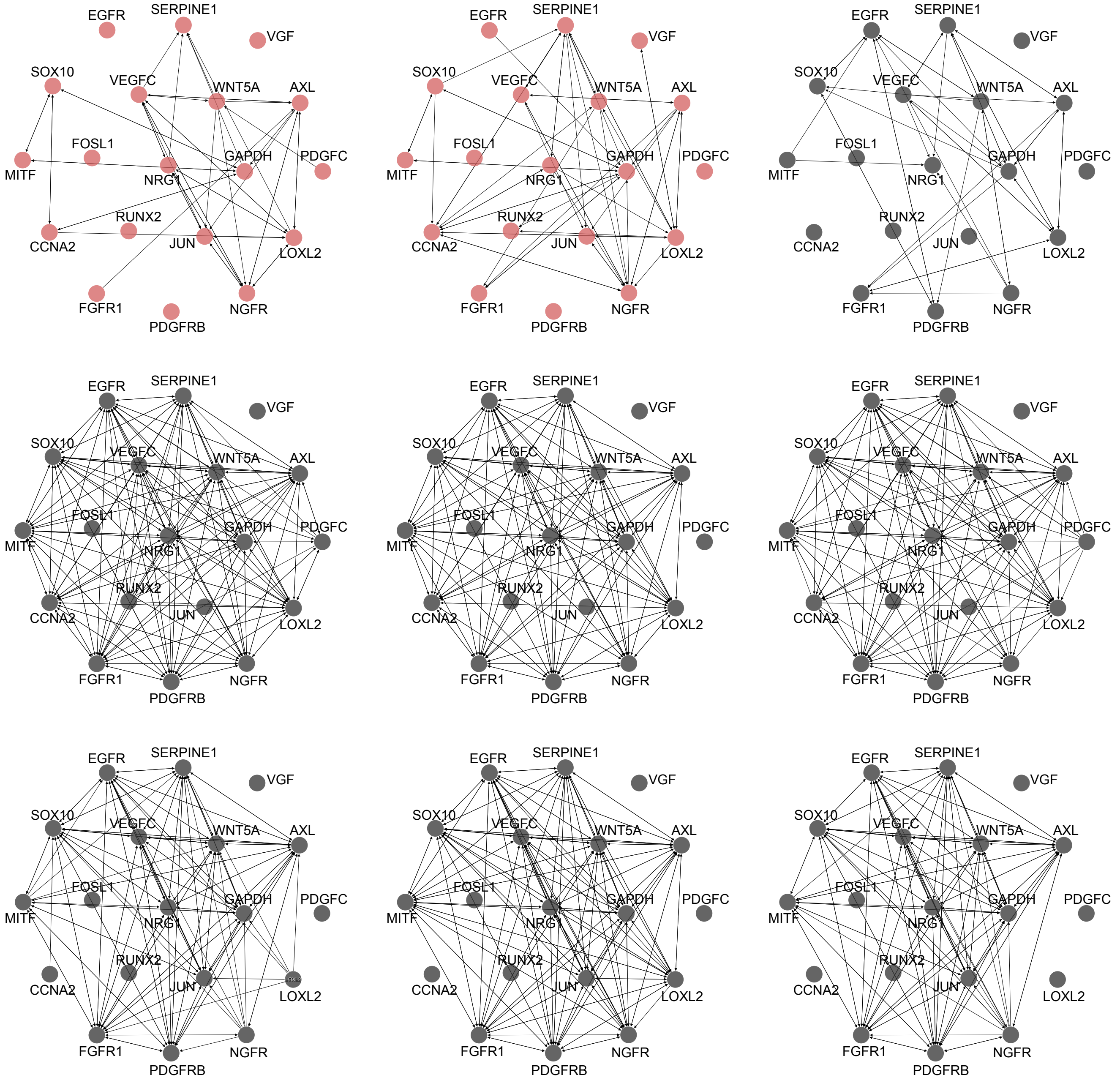


**Figure S8. Related to Figure 4.** The resistant colonies (gray) have more edges in their respective inferred gene regulatory networks than drug naive melanoma cells (red).

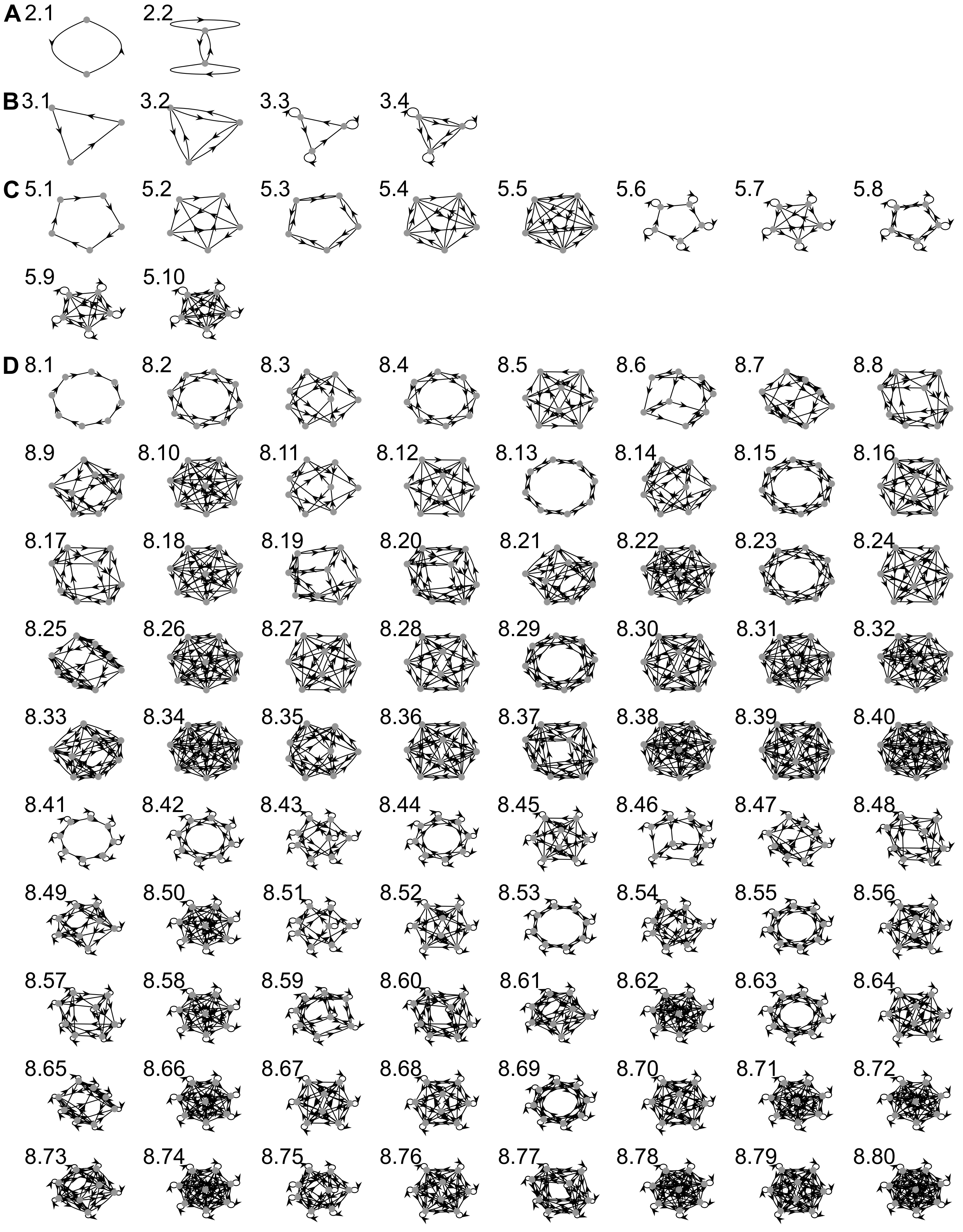


**Figure S9. Related to Star Methods.** All weakly-connected, non-isomorphic, symmetric network architectures of sizes two (A), three (B), five (C) and eight (D).
